# Protoribosomal condensate formation across cationic chemistries

**DOI:** 10.64898/2026.09.29.755306

**Authors:** Simone Codispoti, Valerio G. Giacobelli, Matúš Friček, Václav Verner, Julie Nováková, Martin Mašek, Anton Bokač, Robin Kryštůfek, Radko Souček, Michal Kolář, Giuliano Zanchetta, Klára Hlouchová

**Affiliations:** Department of Medical Biotechnology and Translational Medicine, University of Milano, Segrate, 20054, Italy; Department of Cell Biology, Charles University; Prague, 12843, Czech Republic; Institute of Organic Chemistry and Biochemistry, Czech Academy of Sciences; Prague, 16000, Czech Republic; Department of Physical Chemistry, University of Chemistry and Technology, Prague, 16628, Czech Republic; Institute of Physics of Charles University, Faculty of Mathematics and Physics, Prague, 12116, Czech Republic

**Keywords:** Protoribosome, peptide-RNA coacervation, liquid-liquid phase separation, origins of life, statistical peptide libraries

## Abstract

Early ribonucleoprotein systems likely required mechanisms to concentrate and organize RNA. Modern ribosomes use Mg^2+^ and evolved Lys/Arg-rich protein extensions to stabilize their RNA backbone. Before templated synthesis, however, peptide formation likely generated heterogeneous sequence populations rather than reproducible ribosomal sequences.

Here, we compare Mg^2+^, ribosomal peptides, and statistical peptide libraries in organizing a 136-nucleotide model of the peptidyl transferase center (sPTC). Mg^2+^ condenses sPTC only at a large excess of positive charge following thermal annealing, producing largely arrested libraries. Acidic conditions further promote condensation. Ribosomal peptides instead promote coacervation near charge stoichiometry and form droplets that readily fuse. Several statistical peptide libraries also coacervate with sPTC, despite comprising heterogenous mixtures rather than a single defined sequence. Increasing mean positive charge favors condensation, but Lys/Arg-containing libraries undergo liquid-liquid phase separation more readily and across broader conditions than matched libraries containing the prebiotically plausible diaminopropionic acid (Dpr) and diaminobutyric acid (Dab). Atomistic computer simulations implicate Arg as a major source of this difference, as it can form more numerous and longer-lived hydrogen bonds with RNA, while competitive partitioning experiments show preferential condensate recruitment in the order Arg > Lys > Dab > Dpr.

Together, our findings show that peptide-RNA coacervation can be triggered collectively by statistical peptide ensembles, providing a plausible route to protoribosomal organization without peptide sequence-specific optimization. Cationic chemistry shapes condensate formation and material properties, suggesting that amino acid alphabet formation could have broadened the conditions supporting liquid-like protoribosomal assemblies.

**Significance statement:** How RNA and peptides first became organized into shared compartments is a central question in the origin of life. Using a model of the ribosome’s ancient catalytic RNA core, we show that the chemical identity of positive charge controls not only whether condensation occurs, but also whether the resulting assemblies are arrested or liquid-like. Heterogeneous peptide populations can form RNA-rich droplets without a single defined RNA-binding sequence; however, Lys- and Arg-containing populations do so more readily than populations containing simpler cationic amino acids. Arginine is especially effective, forming persistent contacts with RNA and preferentially entering the condensed phase. Thus, expansion of cationic amino acid chemistry could have broadened the conditions supporting dynamic peptide– RNA compartments during early ribosome evolution.

## Introduction

One of the central challenges in the origins of life research is to understand how the first biopolymers became sufficiently concentrated and organized to interact, polymerize, and ultimately give rise to increasingly complex molecular systems(1–3). Numerous physical mechanisms have been proposed to provide such compartmentalization, including mineral surfaces, lipid vesicles, molecular crowding and phase separation, all of which could have promoted local concentration of reactants and altered reaction kinetics before the onset of biological membranes. However, most models have primarily focused on nucleic acids, whereas early peptides were likely present alongside RNA well before the emergence of modern translation.(4, 5)

The modern ribosome provides perhaps the clearest molecular relic of this transition. Its peptidyl transferase center (PTC), responsible for peptide bond formation, is an RNA-dominated catalytic core that is universally conserved across life and widely regarded as one of the oldest surviving molecular architectures(6–8). Surrounding this ancient RNA core are ribosomal proteins and peptide extensions that are thought to have accumulated during ribosomal evolution(9, 10). An intriguing and largely overlooked feature of the ribosome architecture is the spatial organization of charged species(11, 8). The innermost and evolutionarily oldest regions of the ribosome are largely devoid of proteins. The electrostatic repulsion of the negatively charged RNA backbone is screened primarily by water and coordinated Mg^2+^ ions. In contrast, the outer and more recently accreted layers become enriched in ribosomal proteins containing lysine- and arginine-rich extensions that interact extensively with RNA. This spatial hierarchy raises a fundamental question: how did peptide-borne positive charge add to an RNA core already stabilized by Mg^2+^? RNAs can undergo Mg^2+^-dependent folding(12) and heat-induced phase transitions with lower critical solution temperature behavior but the resulting RNA condensates can be irreversible and dynamically arrested(13). Peptide recruitment may therefore have changed not only the source of charge compensation but also the physical organization and material properties of RNA-rich assemblies.

Peptide-RNA complex coacervation, a form of liquid–liquid phase separation (LLPS), represents one plausible physicochemical mechanism for such reorganization. Such condensates concentrate biomolecules, create distinct chemical microenvironments, and can modulate diffusion and reaction kinetics(14, 15). Recent studies have shown that heterogeneous mixtures of short peptides and oligonucleotides readily undergo phase separation(16, 17). Moreover, both peptide amino-acid composition and sequence influence the physicochemical properties of these droplets(18) and potentially the activity of encapsulated ribozymes(19, 20). We recently showed that mixtures of extant ribosomal peptide fragments spontaneously phase separate with a reconstructed protoribosomal RNA and stabilize its structure, suggesting that peptide-assisted compartmentalization may have preceded the emergence of modern ribosomal proteins(21).

However, reconstructing the earliest stages of ribosome evolution requires moving beyond extant ribosomal proteins, whose compositions have been extensively shaped over billions of years. Prebiotic peptides were instead likely composed predominantly of small aliphatic and acidic amino acids, whereas the canonical basic residues (lysine and arginine) likely became incorporated only later during the evolution of the amino acid alphabet(22, 4, 23). Simpler positively charged amino acids, such as 2,4-diaminobutyric acid (Dab) and 2,3-diaminopropionic acid (Dpr), which were more accessible through prebiotic chemistry, have therefore been proposed as plausible evolutionary intermediates(24– 28, 20).

Here, by using biophysical experiments and computer simulations, we investigate how cationic side-chain chemistry influences peptide–RNA coacervation in a model of early protoribosomal organization. Next to studying individual peptide sequences, we employ randomized (statistical) peptide libraries whose compositions mimic hypothetical stages in the amino acid alphabet evolution. This allows us to differentiate the contribution of composition from that of specific sequences. We find that Mg^2+^ trigger protoribosomal RNA condensation only under restrictive conditions and into largely arrested assemblies, whereas cationic peptides promote dynamic coacervation across a broader phase space. Although increasing net positive charge generally favors condensation, the presence of charge alone does not explain the coacervation propensities: Lys/Arg-containing libraries support LLPS more readily than their Dpr/Dab-containing counterparts, with Arg forming particularly persistent RNA contacts and being preferentially recruited into condensates. These findings suggest that diversification of cationic amino acid chemistry could have expanded and tuned the physicochemical conditions available to emerging peptide–RNA assemblies, even before the evolution of sequence-specific RNA-binding proteins.

## Results

### Mg^2+^ trigger protoribosomal RNA condensation only under restrictive conditions

Throughout the ribosome, the highly negative charge of ribosomal RNA (rRNA) is compensated by a combination of water, inorganic cations and positively charged amino acid side chains. These species are not distributed uniformly across the ribosome(11). Mg^2+^ ions located within 0.5 nm of rRNA are most highly enriched in the immediate vicinity of A2451 (*E. coli* numbering), which serves us as a reference point for the PTC (Fig. 1A). In comparison, Lys and Arg residues are scarce in this inner region and become more abundant at greater distances from the PTC, whereas the negatively charged residues Asp and Glu remain depleted near the RNA throughout the structure (Fig. 1B). The source of positive charge therefore changes radially across the large ribosomal subunit, with Mg^2+^ dominating around the RNA-rich catalytic core, while protein-borne positive charge gets more abundant toward the periphery. In the model of ribosomal accretion, this organization suggests that Mg^2+^ may have supported early PTC rRNA before cationic peptides became increasingly integrated into the emerging ribosome.

**Figure 1.**
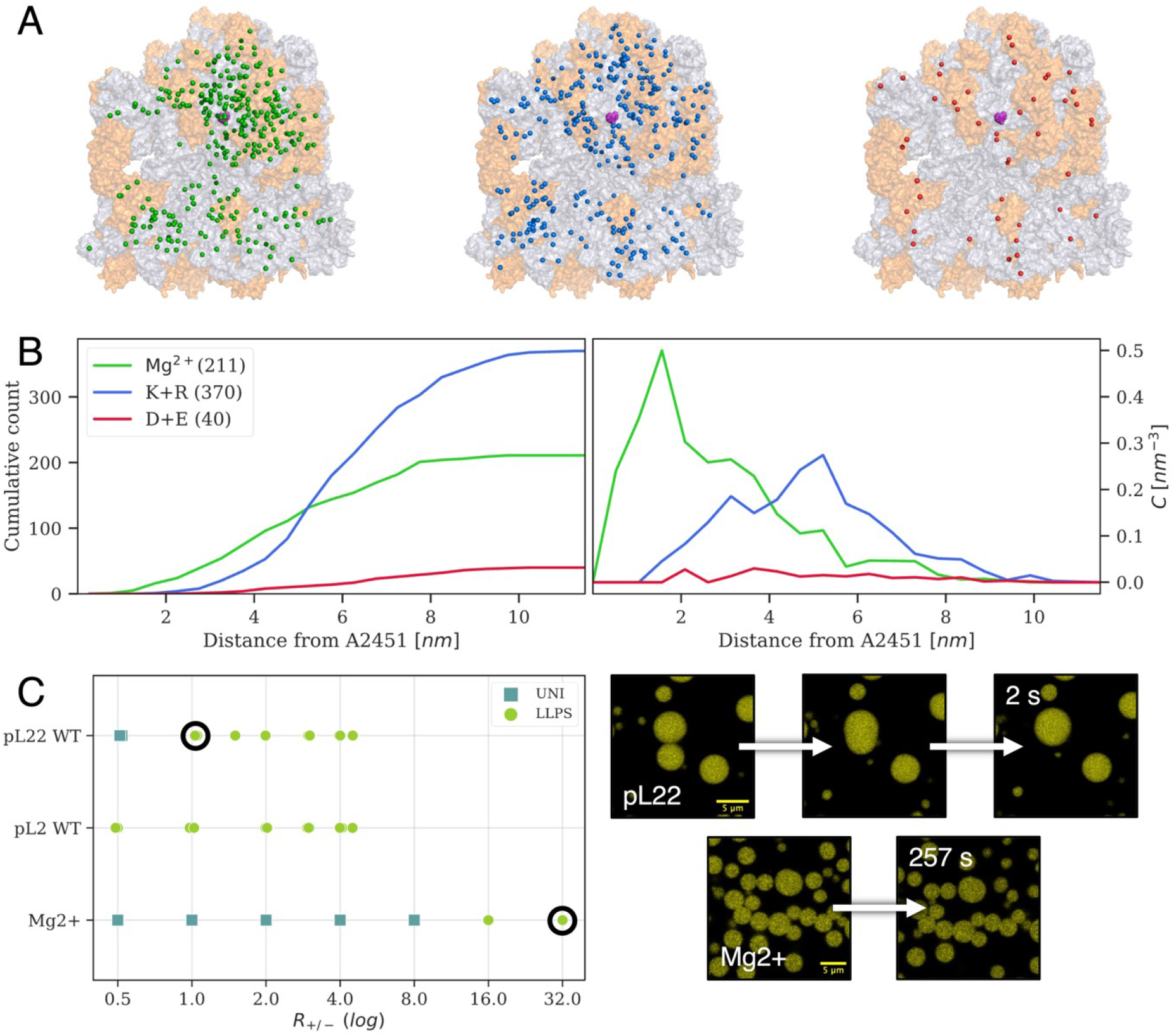
Spatial organization of positive charge in the ribosome and distinct condensation behavior of Mg^2+^ and cationic peptides. (*A*) Spatial distribution of positive charges in the bacterial ribosome (rRNA in gray, rProteins in orange), located within 5 Å from the rRNA. An *E. coli* model with the resolution of 2 Å was used (PDB 7K00). A2451 (shown in magenta) is chosen to represent the center position of the PTC. From left to right, green spheres represent Mg^2+^ ions, blue spheres the alpha carbons of Lys and Arg, red spheres the alpha carbons of Asp and Glu. (*B*) Spatial distribution of Mg^2+^ ions, cationic residues K/R and anionic residues D/E in the ribosomal LSU. On the left, the cumulative distribution as a function of the distance from A2451 is shown; on the right, the same data are represented as the number concentration computed in spherical shells centered on A2451. In parentheses, the total number of counts in the LSU is given. (*C*) Phase behaviour and dynamics of sPTC-Mg^2+^ *vs* sPTC-pL22/2 WT condensates. On the left, a comparison of the LLPS onset (green circles) as a function of the charge ratio R_+/−_ is shown (UNI indicates mixed states, blue squares). The black empty circles highlight the conditions where the confocal time-lapses shown on the right are acquired, for pL22 (upper micrographs) or and Mg^2+^ (lower micrographs) condensates. Both coacervate samples were prepared with the addition of 1x SYBR Green fluorescent dye.

To examine the type of RNA organization that Mg^2+^ alone could support, we used a 136-nt small PTC construct that was studied previously(7, 21) (Table S1, hereafter referred to as sPTC) as a minimal model of protoribosomal RNA. Before mapping its phase behaviour, we tested the stability of sPTC across the experimental conditions used in this study, as Mg^2+^ ions can promote both RNA folding and cleavage(12) (see SI). The RNA remained chemically stable for at least 7 days at Mg^2+^ / RNA charge ratios ranging from R_+/−_ = 0 to 32, both in water and in buffers spanning pH 4-8 (Fig. S1). Here, R_+/−_ denotes the ratio between the total positive charge supplied by the Mg^2+^ and the negative charge of the RNA phosphate groups. We do not have direct experimental access to the detailed sPTC structure in solution; however, consistently with the PDB structure(29), 1D NMR showed a significant base pairing (Fig. S2). Moreover, upon thermal annealing, UV absorbance showed that pairing was largely preserved, with full melting occurring only in presence of large urea concentrations (Fig. S3). Atomistic molecular dynamics (MD) simulations further confirm the overall conformational stability of sPTC at room temperature^21^. Upon heating to 450 K, MD simulations suggest that sPTC remains compact, although the secondary and tertiary structure is notably remodelled. Only about 10% of H-bonds and a few Mg^2+^ ions are lost, while the gyration radius remains stable (Fig. S4), compared to the 300 K reference.

In water, sPTC remained uniformly dispersed at R_+/−_ ≤ 8. Detectable RNA condensates appeared only at a large Mg^2+^ excess, R_+/−_ *≥* 16, and following thermal annealing at 85-95 °C (Figs. S5, S6). Acidic conditions further promoted condensation, with droplets forming at pH 4 following thermal annealing at 95 °C. Mg^2+^ can therefore drive the condensation of sPTC, but only within a narrow range of conditions, involving a large excess of positive charge and favoured by elevated temperature or low pH.

### A ribosomal peptide promotes dynamic RNA coacervation near charge stoichiometry

In our previous study, positively charged ribosomal peptide fragments were shown to induce coacervation of the sPTC construct, with the individual peptide pL22 reproducing the features of the phase behaviour observed for the peptide mixture(21). Here, we used the established pL22-sPTC system as a peptide-mediated reference state and compared it directly with the Mg^2+^ induced condensates described above. pL22 is a positively charged fragment of the extant ribosomal protein uL22 and has the highest proportion (about 44%) of positively charged residues among the ribosomal peptides examined previously (Table S2). While Mg^2+^-induced condensation was detected only at a large excess of positive charge, pL22 WT produces sPTC droplets already close to charge stoichiometry (Figs. 1C, left panel, S7). Similarly, high propensity to LLPS is displayed by another ribosomal peptide, pL2 WT, with fewer positive residues. The ribosomal peptides therefore promote condensation at a substantially lower positive-to-negative charge ratio than Mg^2+^.

The condensates formed by the two positively charged species, Mg^2+^ ions and peptides, also differed markedly in their dynamics. pL22-sPTC droplets readily coalesced, with fusion completed within seconds (Fig. 1C, right panel). By contrast, despite their round morphology, the Mg^2+^-induced condensates did not visibly coalesce over minutes and hours. This suggests that they represent dynamically arrested RNA-rich structures rather than fully liquid coacervates. Interestingly, the appearance of irreversible, percolated droplets upon thermal annealing has been recently observed at similar Mg^2+^ excess for RNA strands with high purine content(13), close to the composition of PTC, while strands with lower purine content retained fluidity. Peptide-mediated condensation therefore did not only shift the phase boundary, it changed the properties of the condensed phase from arrested to a liquid-like coacervate.

These two modes of condensation offer an experimental model for the transition proposed by the accreted distribution of charge within the ribosome. Mg^2+^ can promote condensation of the PTC RNA model, but only under restrictive conditions and into a largely arrested state. Ribosomal peptides support condensation under broad conditions while permitting rearrangements of the condensed phase. Recruitment of cationic peptides during ribosomal evolution could therefore have contributed to widen the range of conditions compatible with protoribosomal RNA compartmentalization and their dynamics. At the same time, pL22 is an evolved sequence in which net charge, charge distribution, amino acid composition and sequence are all coupled. We therefore next asked whether any cationic peptide with a sufficient net charge could support liquid coacervation and whether the chemical identity of the positively charged side chains also determines the phase behaviour.

### Peptide libraries containing prebiotically plausible basic amino acids display LLPS

To distinguish the general contribution of peptide charge from the effects of individual amino acid chemistries and sequences, we examined sPTC phase behaviour in the presence of various statistical peptide libraries. Their compositions were built around a proposed early set of ten proteinogenic amino acids - Ala, Asp, Glu, Gly, Ile, Leu, Pro, Ser, Thr and Val - interred from their occurrence and relative abundance in meteorites and prebiotic synthesis experiments(30). This set contains the acidic residues Asp and Glu but no basic amino acids, and therefore lacks sidechain positive charge that could support association with RNA. We used the eight uncharged members of this set - Ala, Gly, Leu, Ile, Pro, Ser, Thr, and Val (denoted as 8E) - as a common compositional background and systematically varied the amount and chemical identity of the charged residues. As models of simpler cationic chemistry, we included Dpr (hereafter O) and Dab (hereafter U), short-chain diamino acids proposed as prebiotically plausible alternatives for the canonical basic residues Lys and Arg(20, 24, 25, 27, 28) (Fig. 2A). Each library comprised a heterogenous population of amino acids peptide sequences produced using isokinetic mixture solid-phase synthesis, from a defined bulk amino acid composition(31) (Fig. S8). A library containing 8E together with Asp and Glu (8E+ED; mean net charge Q_net_ = −5) served as an acidic reference corresponding to the proposed early proteinogenic set. Six further libraries were organized in three pairs with designed mean net charges of Q_net_ = 0, +5, and +7. Within each pair, a Dpr/Dab-based library was compared with its Lys/Arg-based counterpart: 8E+UOED with 8E+KRED, 8E+UO with 8E+KR, and ribUO with ribKR (Fig. 2C). The latter pair was designed to match the net charge of the ribosome pL22 WT peptide (hence the “rib” resignation). Dpr and Dab differ from Lys and Arg in both side-chain length and basicity, with approximate pKa values of 8.2 compared with 10.5 for Lys and 12.5 for Arg (Fig. 2A). This design allowed us to examine compositional effects without relying on the behaviour of a single evolved peptide, while exploring the behavior of populations with a relatively broad charge distribution (Fig. S9).

**Figure 2.**
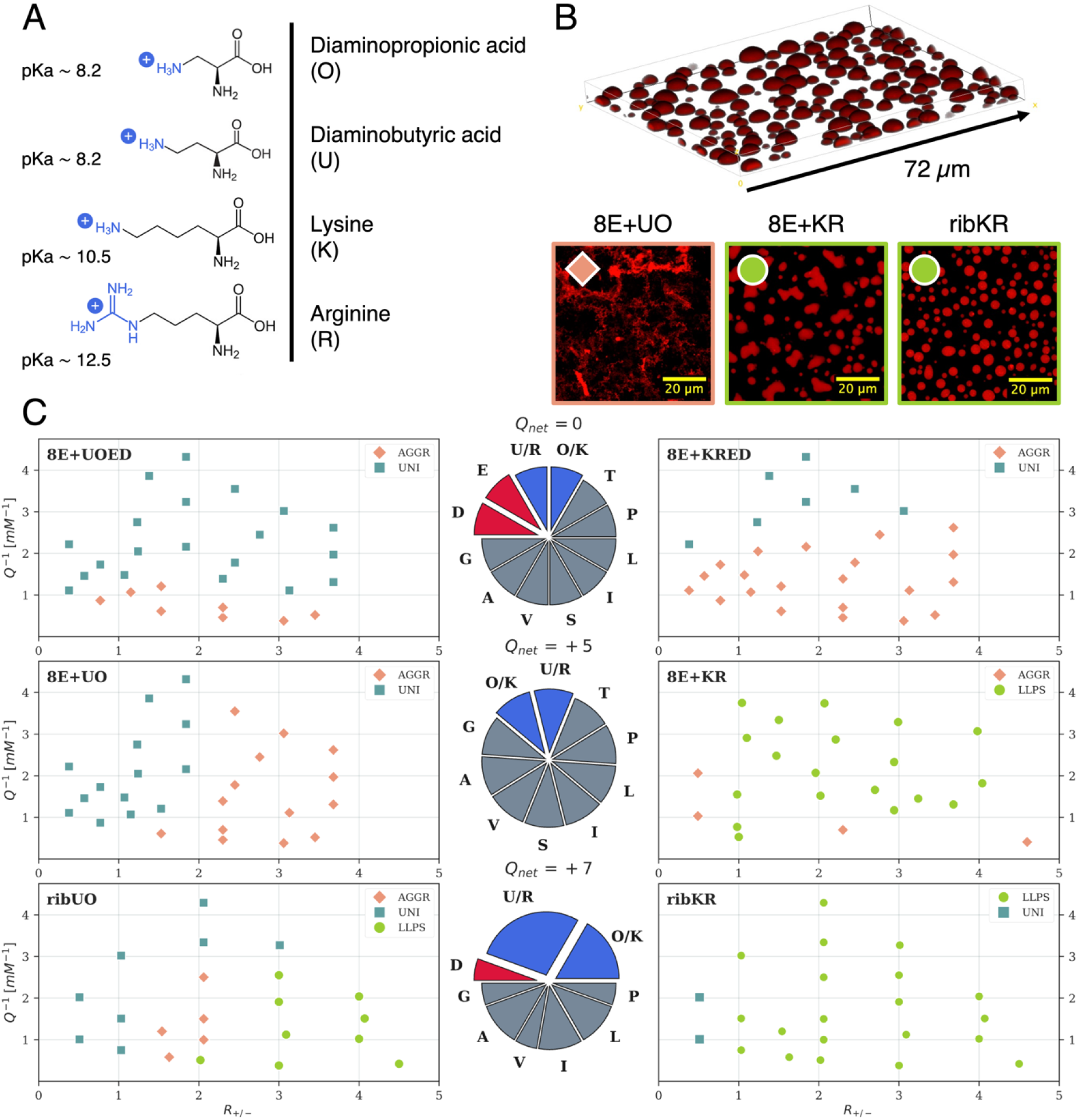
Cationic side-chain chemistry shapes peptide–RNA condensation and phase behavior. (*A*) Chemical structures of the canonical cationic amino acids Arginine (R), Lysine (K) and of the noncanonical, shorter versions of Lysine: Diaminobutyric acid (U) and Diaminoproprionic acid (O). The pKa of the basic groups, highlighted in blue, is also reported. (*B*) On the top, confocal 3D-slab reconstruction of sPTC-pL22 WT condensates at charge ratio R_+/−_ = 1; on the bottom, representative micrographs showing aggregation in sPTC-8E+UO mixtures (R_+/−_ = 3) and condensates with different morphologies in sPTC-8E+KR and sPTC-ribKR mixtures (R_+/−_ = 1). Fluorescent samples were obtained with the addition of 500 nM TAMRA-labeled peptides. (*C*) Experimental phase diagrams of “early” combinatorial libraries (left column) and of their Arg/Lys-containing counterparts (right column), as a function of the charge ratio R_+/−_ and of the inverse of the total concentration of charged units Q^-1^. Blue squares indicate mixed states (UNI), red diamonds the presence of aggregates (AGGR) and green circles the formation of liquid-like droplets (LLPS). On the center, pie-charts display the composition of the employed libraries, with acidic/basic residues highlighted in red/blue.

As expected, the acidic 8E+ED library did not interact detectably with the negatively charged RNA in solution and did not display phase separation under any of the explored conditions. Neither of the two net-neutral libraries (8E+UOED and 8E+KRED) underwent LLPS over the tested range of concentrations and peptide-to-RNA ratios. The samples remained uniform or formed aggregates at high total concentration (Fig. 2B, C). Instead, increasing the designed mean net charge to +5 produced different phase behaviour in the two residue sets. The Dpr/Dab-containing 8E+UO library remained uniform or formed aggregates throughout the sampled phase space, while the corresponding Lys/Arg-containing 8E+KR library formed droplets already at room temperature and over a broad range of conditions (Fig. 2C). Although some of the droplets displayed irregular shapes (Figs. 2B, S10), their inner fluidity was tested through FRAP experiments, in which the recovery of the fluorescent signal provides information about the mobility of molecules in a bleached portion of the droplet and in its surrounding regions(32, 33) (Fig. S11). The low percentage of recovery may indicate the coexistence of mobile and arrested fractions within the sample.

At Q_net_ = +7, both libraries triggered liquid droplet formation. However, for the Dpr/Dab-containing ribUO library LLPS remained restricted to a subset of the tested conditions and was accompanied by uniform and aggregated states elsewhere in the phase diagram (Fig. 2C). The Lys/Arg-containing ribKR library, instead, formed droplets across nearly the entire examined range. Their shape and degree of recovery revealed a fully liquid behaviour (Figs. 2B, S10, S11). Increasing net positive charge therefore enhanced the propensity of both residue sets to condense sPTC, but the Dpr/Dab-containing libraries required a greater cationic content and retained a substantially narrower LLPS regime than their Lys/Arg-containing counterparts.

Our results suggest that, while higher mean net charge overall enhanced coacervation propensity, the type of cationic residues played a major role in determining the phase behaviour.

### The binding mode of arginine with RNA is a key determinant of condensate properties

To identify the molecular interactions underlying the different phase behaviour of the ribKR and ribUO libraries, we performed atomistic MD simulations of sPTC in the presence of sequence-defined peptide representatives. The statistical libraries used in the experiments comprise heterogeneous populations of peptides that differ in their individual sequences. Explicit representations of this heterogeneity at atomistic resolution would require sampling a huge number of potential library members and would complicate the assignment of RNA contacts to individual side chain chemistries. We therefore selected two pL22-derived peptides as sequence-defined proxies for the corresponding libraries.

pL22 WT clearly matches the net charge and composition of the ribKR library, while pL22 UO (with Dpr/Dab substituting Lys/Arg) is a proxy of the ribUO library. Importantly, the experimentally determined phase diagrams of pL22 WT and pL22 UO mixtures with sPTC closely reproduce those of ribKR and ribUO, respectively (Fig. S7). The simulations show that both types of peptides readily bind the RNA, replacing only a few of the bound Mg^2+^ ions (Figs. 3A, S12, S13). As a result of mostly electrostatic interactions, the peptides cover the sPTC surface uniformly, with no detectable specificity (Fig. S13). In both cases, the tertiary structure of RNA is largely preserved, with stable secondary structure and slight compaction upon binding (Fig. S14)(34, 21).

**Figure 3.**
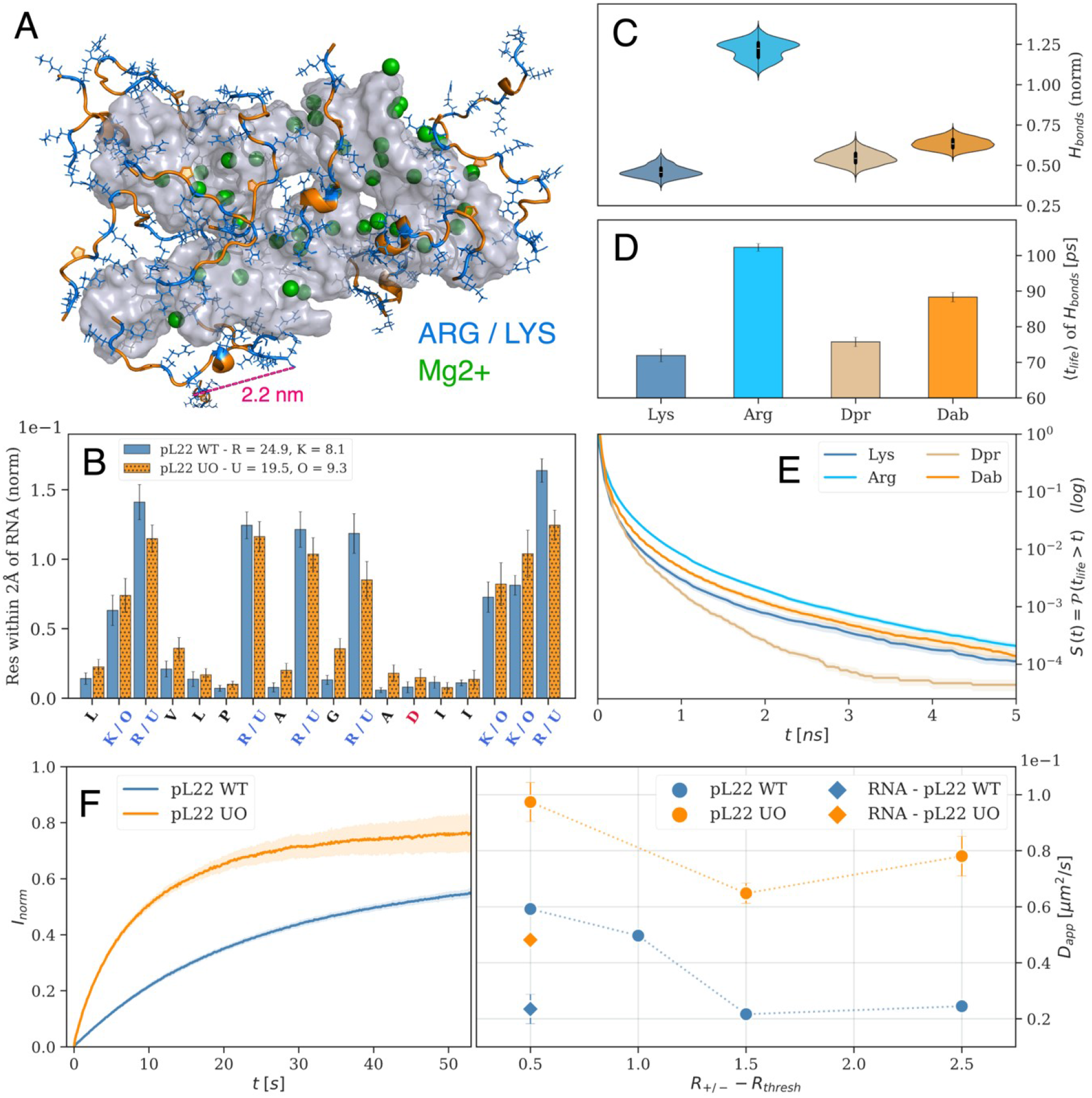
Cationic side-chain chemistry modulates peptide–RNA interactions and condensate dynamics. (*A*) Representative simulation snapshot of an equilibrated bound-state of a sPTC-pL22 WT mixture. The RNA surface is shown in gray, Mg^2+^ ions are represented by green spheres, and peptide backbones are highlighted in orange, while their Arg/Lys are colored in blue. (*B*) Average fraction of pL22 WT (blue, filled) or pL22 UO (hatched, orange) residues within 2 Å from RNA. Per-residue counts are normalised by the total counts, with the legend showing non-normalized averages for single residues. Error bars represent standard errors of the means over the 10 replicas. (*C*) Probability density estimates of H-bonds between RNA and Arg, Lys, Dab or Dpr. The number of H-bonds is normalized by the number of bound peptides in each replica and by the occurrence of each residue in the peptide sequence. The cumulated time-averaged values of 10 replicas are shown. (*D-E*) Average lifetime of H-bonds (*D*) between RNA and Arg, Lys, Dab or Dpr, as estimated by the integral of the average survival functions 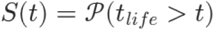 (*E*). (*F*) On the left, averages and standard deviations of normalized FRAP signals from pL22 WT (blue) or pL22 UO (orange) condensates (n=6) at R_+/−_ = 3. On the right, apparent diffusion coefficients estimated from fitting normalized FRAP signals with an exponential recovery, as a function of the charge ratio R_+/−_ shifted by the approximate threshold value R_thresh_ for the onset of LLPS. Circles represent diffusion coefficients of peptides, while diamonds are related to estimates of sPTC diffusion from FRAP experiments of SYBR Green stained condensates.

However, when closely inspecting the local interactions between amino acids and nucleotides, differences between canonical and noncanonical amino acids emerge. Fig. 3B shows the average number of contacts experienced by each peptide residue with the RNA. In pL22 WT, the positively charged residues clearly stand out from the other amino acids, with Arg in particular remaining in closer contact with RNA. Instead, pL22 UO forms more homogeneously distributed contacts among all residues, suggesting weaker, more dynamic interactions. The stronger binding of WT peptides with sPTC is also reflected by their larger binding interface, inaccessible to solvent molecules, which is proportional to the binding free energy (Fig. S12). The origin of such diversity, irrespective of the adopted peptide force field (Fig. S15), can be mostly attributed to the fact that Arg can form a larger number of H-bonds with RNA than both Lys and the noncanonical residues (Fig. 3C), with longer lifetimes (Fig. 3D, E). The overall picture emerging from the simulations is that of Arg interactions dominating over the other residues.

Consistently, the experimental mobility of peptides and RNA inside coacervate droplets, estimated from FRAP and expressed through the apparent diffusion coefficient in Fig. 3F, is higher for pL22 UO than for pl22 WT at all distances from the phase boundary, corresponding to faster rearrangements of peptide-RNA contacts at the molecular scale.

### Selective partitioning of cationic peptides into coacervates scales with side-chain length and cationic group identity

To further probe how peptide side-chain chemistry governs their recruitment into RNA coacervates, we performed a competitive partitioning assay. sPTC was mixed with an equimolar mixture of either the pL22 WT and pL22 UO peptides, or the ribKR and ribUO libraries at R_+/*−*_ = 3, a condition that triggered LLPS with each sample separately. Differences in the uptake of basic amino acids were assessed by amino acid analysis of the initial solution and of the dilute phase (Fig. 4A). Interestingly, pL22 showed a higher level of depletion than the library, yet in both setups, selective depletion of the canonical basic residues from the dilute phase was observed (Fig. 4B). This ranking reflects preferential uptake of longer side chains (Lys, Arg vs. Dab, Dpr) into the condensed phase. Notably, no significant difference between Arg and Lys was observed in the ribKR/UO sample. Small differences between Arg and Lys, or between Dab and Dpr, likely reflect limitations of the experimental setup rather than actual differences, since the uptake of both residues within a pair should be identical in the pL22 WT/UO setup.

**Figure 4.**
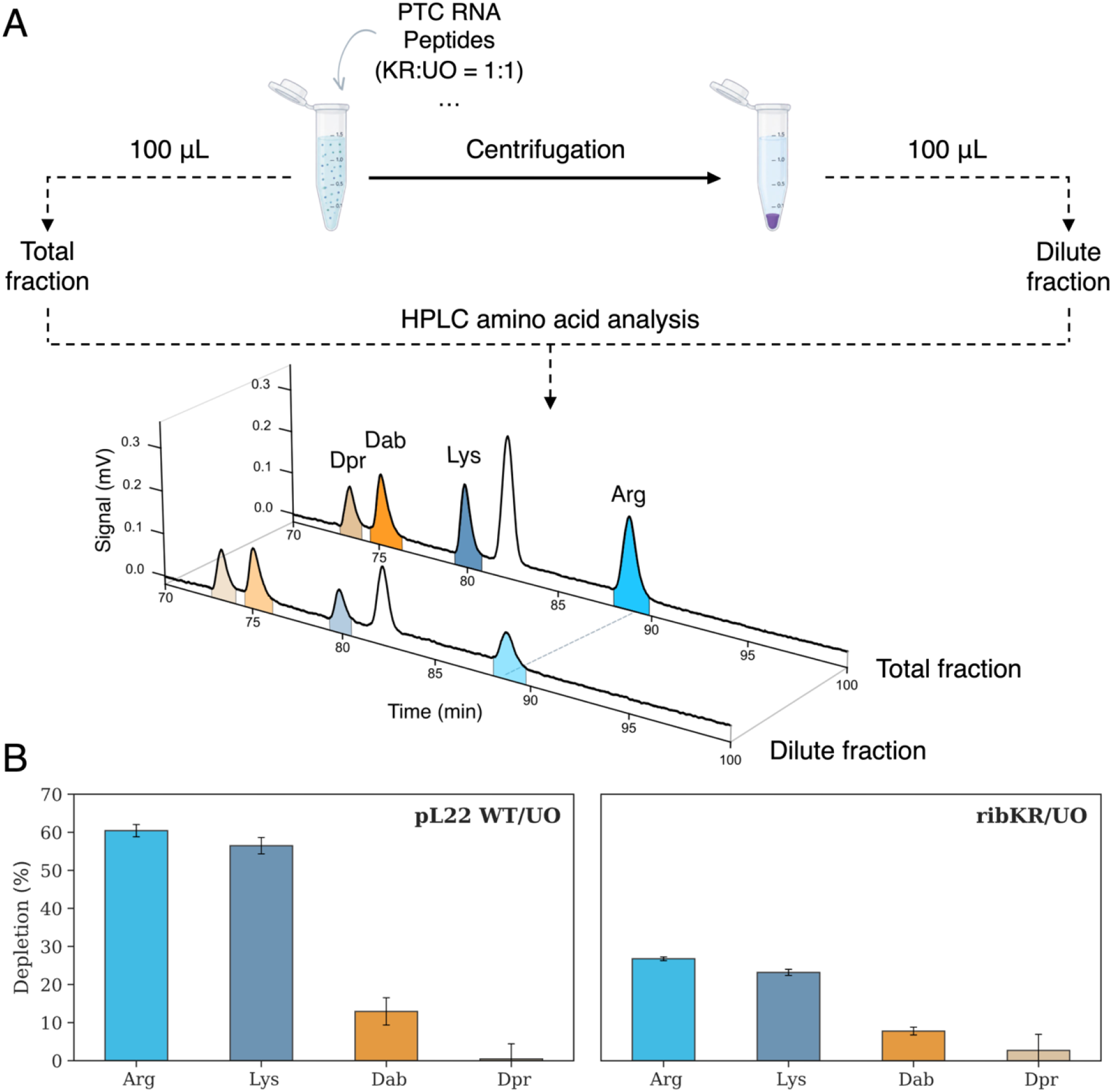
Competitive partitioning of basic amino acid residues into RNA-rich condensates. (A) Schematic representation of the competitive partitioning assay. A 1:1 mixture of the WT and UO variants of pL22 (or of ribKR and UO) was mixed with protoribosomal RNA at overall charge ratio R_+/−_ = 3 to induce liquid–liquid phase separation. Total fraction and dilute fraction (100 µL) were collected, acid-hydrolyzed, and subjected to amino acid analysis by HPLC. The chromatograms show representative amino acid profiles of the total and dilute fractions; the reduced peak areas of the basic amino acids in the dilute fraction are visible. (B) Quantification of the depletion of basic amino acids from the dilute phase for pL22 WT/UO (left) and ribKR/UO (right). Arg, Lys, and the basic analogs Dap and Dab were quantified in the total and dilute fractions and are expressed relative to the total fraction (mean ± SE, n = 5). Preferential depletion of a residue from the dilute phase indicates partitioning of the corresponding variant into the condensed, RNA-rich phase.

## Discussion

The modern ribosome contains an RNA core that is widely regarded as its evolutionary oldest region(6, 7). Mg^2+^ ions and water compensate for much of the negative charge in this core, while positively charged protein segments become more abundant toward the ribosomal periphery. This organization raises a simple question: what could peptide-borne positive charge have added to an RNA system already supported by metal ions? Beyond charge compensation itself, peptides may have changed the physical and catalytic properties of the growing protoribosomal structure and the conditions under which the complex may have been integrated into separate compartments.

Such concentration and physical reorganization could arise through peptide-mediated coacervation. Positively charged, unstructured fragments of ribosomal proteins that extend toward the PTC, including pL22, were previously shown to form coacervates with PTC model RNAs and to enhance their structural stability(21). However, these extant peptides contain Lys and Arg, whereas earlier peptide synthesis may initially have drawn on a broader set of cationic building blocks, including the prebiotically plausible diamino acids Dpr and Dab, irrespective of whether these residues ever became part of a genetically encoded amino acid repertoire. This raises the question of how the availability of different cationic side chains could change the ability of peptide populations to recruit RNA and shape the properties of the resulting condensed phases. Here, we use sPTC condensation as a readout of how the amount, distribution, and chemical identity of positive charges could have shaped early peptide–RNA organization.

We show that Mg^2+^ and peptide mediated charge produced different forms of sPTC organization. Mg^2+^ alone condensed sPTC only at a large excess of positive charge and under conditions favored by thermal annealing or low pH. The resulting assemblies showed no coalescence and appeared largely arrested. By contrast, pL22 promoted condensation close to charge stoichiometry and produced droplets that readily coalesced. Presenting multiple, distributed positive charges on a flexible peptide therefore expanded the conditions compatible with sPTC condensation and changed the material state of the resulting RNA-rich phase, which was previously shown to favour catalytic activity(13).

The statistical peptide libraries extend our comparison from a single, through evolution highly optimized ribosomal sequence to more prebiotically plausible heterogeneous peptide populations. Before templated synthesis, peptide formation would not have repeatedly generated a single sequence such as pL22 with high fidelity. It would instead have sampled a broad sequence space shaped by the available amino acids, the chemical selectivity of polymerization or other so far unknown mechanisms. Experimental models of prebiotic peptide synthesis similarly produce highly diverse sequence populations rather than one predominant product(35). Several of the statistical libraries examined here nevertheless underwent LLPS with the sPTC. Thus, peptide–RNA coacervation can emerge as a population-level property of heterogeneous sequence space.

At this population level, mean net charge differs from the charge of a sequence-defined peptide. A library with a mean charge of zero contains cationic, anionic, and mixed-charge peptides, while net-positive libraries likewise span a distribution of individual charges. Increasing the mean positive charge shifts this distribution toward a larger fraction of peptides capable of multivalent RNA interaction. Indeed, it increases the number of sequences containing local clusters of positive residues, as captured by Sequence Charge Decoration(36) values, and the typical length of such clusters (Fig. S9). The broader coacervation window of the ribKR and ribUO libraries may therefore reflect both their greater mean charge and the increased abundance of highly cationic patches, which can contact several RNA groups simultaneously. This is consistent with the recently reported behaviour of short homopeptides, whose LLPS propensity increased with increasing length of Lys and Arg stretches. However, too many consecutive charged residues may lead to substitution of Mg^2+^ ions(34), unfolding of RNA and formation of solid precipitates(33, 37). Phase behavior is thus determined by the distribution of sequence properties across the library rather than only by mean composition alone.

The comparison between compositionally related libraries further shows that net positive charge is necessary but insufficient to determine the condensate properties. At the same mean charge, Lys/Arg-containing populations accessed LLPS more readily and across a broader range of conditions than the corresponding Dpr/Dab-containing populations. Dpr and Dab were not intrinsically unable to support coacervation, because the ribUO library formed liquid droplets at sufficiently high mean positive charge. Their incorporation instead produced a narrower LLPS profile and a greater tendency toward aggregation. The chemical identity of the cationic side chains, their conformational degrees of freedom and binding modes therefore influence whether and how efficiently a peptide population converts electrostatic interactions with RNA into a liquid condensed state. In sequence-defined systems, it was indeed found that peptides containing Arg formed coacervates with polyU with larger droplets and at lower peptide concentrations than peptides containing Ornithine, a simpler basic amino acid(27). Analogously, oligomers of Orn, Dab or Lys yielded coacervation – and enhancement of the activity of a ribozyme – with different effectiveness(38).

The atomistic MD simulations identified Arg as a major source of the difference between these cationic chemistries. Both pL22 WT and pL22 UO associated broadly with the sPTC surface, showing that Dpr/Dab substitution did not prevent electrostatic RNA binding. However, in wild-type pL22 Arg formed more H-bonds, longer-lived contacts, and a larger binding interface with RNA. The prominent role of Arg agrees with its widespread presence in contemporary RNA-binding proteins. Its guanidinium group can form multiple hydrogen bonds with neighboring RNA groups, and Arg-rich peptides commonly bind RNA more strongly than Lys-rich analogues(39–41). Arg can also generate different condensate phase boundaries and material properties(42–44). In the present system, Arg therefore provides modes of RNA interaction beyond electrostatic attraction alone.

This interaction difference was also observed when the ribKR vs. ribUO libraries and pL22 WT vs. pL22 UO peptides competed for recruitment into the same sPTC condensates. Amino acid analysis of the dilute phase showed preferential depletion in the order Arg>Lys>Dab>Dpr, indicating that peptides containing canonical basic residues (Arg in particular) were more strongly recruited into the condensed phase. Because the analysis reports the composition of the partitioned peptide populations rather than the behaviour of free amino acids, this ranking reflects selection among the peptide sequences. The greater recruitment of Lys and Dab or Dpr is compatible with contributions from side chain reach and basicity, while the preference for Arg over Lys points to the additional capacity of the guanidium group. Such differential partitioning could provide a physicochemical route for enriching specific peptide chemistries before the emergence of templated RNA-binding proteins(45, 46).

Whether peptide–RNA condensation contributed to ribosome evolution remains unknown(47). If it did, coacervation could have prevented dilution and increased the local concentrations of RNA, peptides, substrates, and cofactors. These effects might have favored otherwise inefficient reactions, but concentration alone does not imply catalytic benefit. Strong interactions may also restrict molecular exchange, trap nonproductive RNA conformations, or inhibit product release(34, 13, 48, 49, 20). Moreover, the available structural measurements indicate that sPTC is substantially folded before condensation, but its detailed organization within droplets remains to be determined(50). Direct measurements of sPTC structure and activity within Mg^2+^-, Dpr/Dab-, and Lys/Arg-mediated condensed states are therefore essential.

Our results do not necessarily establish LLPS as an intermediate in ribosome evolution or as a driver of amino acid alphabet expansion. They show that the amount, distribution, and chemical identity of peptide-borne positive charge strongly alter the organization of a model protoribosomal RNA. Statistical peptide libraries further demonstrate that coacervation can emerge from heterogeneous sequence populations without prior specification of an evolved RNA-binding peptide. If RNA-rich condensed phases contributed to early ribonucleoprotein systems, their ability to oppose dilution and preferentially enrich specific peptide chemistries could have generated compositional biases toward more effective RNA binders, with Arg providing a particularly consequential expansion of the available interaction repertoire.

## Materials and Methods

### Synthesis of sPTC RNA

The sPTC RNA construct (Table S1) was synthesized according to the procedure reported in **(21)** with minor modifications. For more details, see SI.

### Synthesis of individual peptides and peptide libraries

WT and UO versions of pL2 and pl22 (Table S2) were synthesized as described in (21), using standard protocols for solid-phase peptide synthesis. The identities and purities of the peptides were confirmed by mass spectrometry using UltrafleXtreme MALDI-TOF/TOF mass spectrometer (Bruker Daltonics, Bremen, Germany) according to the standard procedure. Peptide libraries were synthesized using the isokinetic synthesis method(51), with coupling mixtures of amino acids whose quantity was adjusted according to their respective reactivities. The length of the peptides within the libraries varied between 18 (for ribKR and ribUO) and 25 (for 8E+KR, 8E+UO, 8E+ED, 8E+KRED, 8E+UOED) and N-end fluorescent labelling was used for selected samples. For more details, see SI.

### NMR spectroscopy

NMR experiments were performed on a Bruker Avance III HD 850 MHz spectrometer equipped with a 5 mm inverse-detection probehead (Z131194_0003) and operated using TopSpin 3.6.5 (Bruker) on CentOS 7.4. RNA samples were prepared at a concentration of 60 μM in H2O/D2O (9:1, v/v) in 3 mm NMR tubes, with a final sample volume of 160 μL. Spectra (Fig. S2) were acquired at 298 K, with temperature maintained using the instrument’s variable-temperature unit. Water suppression was achieved using the standard Bruker ZGESGP pulse sequence, employing excitation sculpting with frequency-selective shaped pulses and pulsed-field gradients.

### Microscopy Analysis of Mg^2+^-Induced Phase Separation

Phase separation of 8.4 μM sPTC RNA was evaluated in the presence of Mg^2+^ at charge ratios R_+/−_ of 0, 4, 8, 16, and 32 upon thermal annealing at different pH values. Following thermal cycling, all samples were incubated at RT and analyzed at 24 h and 7 d intervals in 96-well flat-bottom Costar plates using a Leica DMi8 inverted microscope equipped with a 40× objective, in bright-field and Differential Interference Contrast (DIC). See SI for more details.

### Coacervation

Phase separation diagrams were estimated as a function of the R_+/−_ ratio and of the molar charge concentration (in units of the elementary charge) Q of the mixtures defined as:

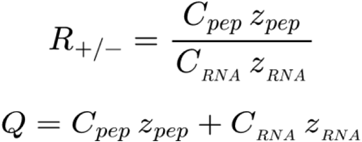

where *C*_*pep*_ and *C*_*RNA*_ are the molar concentrations of peptides/peptide libraries and rRNA, respectively, and *z*_*pep*_, *z*_*RNA*_ their charge numbers. The R_+/−_ ratio represents the stoichiometric charge ratio of the molecules in solution (positive over negative charges), while Q denotes the molar concentration of charged units. Details about the charges of individual components, sample preparation and microscopy characterization are provided in SI.

### Confocal Imaging and FRAP measurements

Samples were prepared as for phase diagram characterization, with the addition of: (i) 500 nM TAMRA-labeled peptides/peptide library for sPTC-peptides coacervates; or (ii) 1x SYBR Green fluorescent dye for sPTC-Mg^2+^ condensates. Confocal imaging and Fluorescence Recovery After Photobleaching (FRAP) experiments were performed on a WLL scanning Confocal Microscope (Leica Stellaris 8) equipped with a 63× (NA = 1.40) oil immersion objective (HC PL APO CS2) and a sCMOS camera Leica DFC9000 GTC. The imaging system was operated using the LASX (LEICA) Software. More details about sample preparation, photobleaching procedure, image acquisition, data treatment and analysis are given in SI.

### Competitive partitioning assay

Samples for peptide partitioning determination were prepared by mixing 1.6 µM sPTC RNA, 1 mM MgCl2, 20mM Tris (pH 7.5). LLPS was induced by adding 96 uM peptide mixture (1:1 equimolar ratio of pl22 WT/UO or ribKR/UO library). Samples were left at room temperature for 30 min and 100 µL were transferred to the new tube (Total fraction). The rest of the sample was centrifuged at 20 000×*g* for 20 min at 20 °C. 100 µL of clarified sample (Dilute fraction) were transferred to the new tube. Samples were prepared in pentaplicates and submitted for amino acid analysis (see SI).

### Simulations of bare RNA

The sPTC structure with bound Mg^2+^ ions was extracted from the experimental model of the extant *E. coli* ribosome (PDB 5AFI(29)) as described previously(21). The sPTC was placed in a periodic rhombic dodecahedral box with a minimum solute-box distance of 1.3 nm. The box was solvated with water and neutralized with a sufficient number of K^+^ ions, followed by the random addition of excess 150 mM KCl and 20 mM MgCl2. In total, 60 Mg^2+^ ions were present in the box, including both the structural ions (initially present in the PDB model) and the excess ions added during system preparation. The RNA was described using the ff10 force field together with the OPC water model and the corresponding ion parameters. For more details about the simulation conditions, equilibration protocol and data analysis, see SI.

### Simulations of RNA-peptide mixtures

The RNA-peptide mixture was modeled using a periodic rhombic dodecahedron box with a volume of 5000 nm^3^ containing one sPTC and 20 peptide molecules, roughly corresponding to a charge ratio R_+/−_ of 1. Twenty copies of the peptide (obtained clustering of the single peptide simulations) were inserted into the box at random positions and orientations, yielding 10 independent initial configurations. Each box was subsequently solvated with SPC/E water, neutralized with chloride ions, and supplemented with 10 mM excess MgCl_2_. Details about equilibration protocols and simulation conditions, see SI. Trajectories were analyzed using a combination of GROMACS analysis tools and custom *Python* scripts. Measured quantities were either averaged among the 10 different replicas and errors estimated as standard error of the mean or cumulated together. See SI for the description of the simulation analysis, including RNA-peptide contacts, H-bonds lifetime, binding interface, distribution of cations and RNA structural analysis.

## Acknowledgements

We thank Oliver Hartley for providing some of the labelled peptides, Pavel Srb for 1D NMR, Andrea Spitaleri and Kosuke Fujishima for discussions.

The work was supported by the Czech Science Foundation (project 25-15428S) and 4EU+ (SEED4EU+ RECHARGE and visiting professorship to G.Z.). We also acknowledge VSB – Technical University of Ostrava, IT4Innovations National Supercomputing Center, Czech Republic, for awarding this project access to the LUMI supercomputer, owned by the EuroHPC Joint Undertaking, hosted by CSC (Finland) and the LUMI consortium through the Ministry of Education, Youth and Sports of the Czech Republic through the e-INFRA CZ (grant ID: 90254).

The use of the confocal microscope was supported by the Department of Excellence Project “SCALE UP” funded by the Italian Ministry of University and Research (MUR), Department of Medical Biotechnology and Translational Medicine. Finally, we acknowledge ISCRA for awarding this project access to the LEONARDO supercomputer (grant ID: HP10CHE51C), owned by the EuroHPC Joint Undertaking, hosted by CINECA (Italy).

## Author Contributions

Conceptualization: KH, MK, GZ. Methodology: KH, MK, GZ, SC, VGG, MF, VV, JN, MM, RK, RS, AB. Investigation: KH, MK, GZ, SC, VGG, MF, VV. Visualization: VGG, SC, AB, MK. Funding acquisition: KH, MK, GZ. Project administration: KH, MK, GZ. Supervision: KH, MK, GZ, SC, VGG. Writing – original draft: KH, MK, GZ, SC, VGG, MF, VV, AB.

