## Supplementary material for "Protoribosomal condensate formation across cationic chemistries": Manuscript_Supplementary Inofmration Biox.pdf

**This PDF file includes:**

Supporting text
Figures S1 to S15
Tables S1 to S2
References

### Supplementary Text

#### Synthesis of sPTC RNA

To synthesize the sPTC RNA construct, a double-stranded DNA (dsDNA) template was assembled by annealing equimolar amounts (15 pmol each; 8.1 µg total DNA) of sPTC_F and sPTC_R single-stranded DNA oligonucleotides purchased from Sigma-Aldrich (Table S1) in NEBuffer 2 to a total volume of 47 µL. The mixture was denatured at 80 °C for 5 min and cooled to room temperature at 0.1 °C $s^{-1}$. Second-strand synthesis was performed by adding 5 U of Klenow Fragment of DNA polymerase I (NEB) and 2 µL of 10 mM dNTP mix (SERVA), followed by incubation at 25 °C for 1 h and thermal inactivation at 50 °C for 15 min. The resulting dsDNA template was purified using the Monarch PCR & DNA Cleanup Kit (NEB). *In vitro* transcription was conducted with the HiScribe T7 High Yield RNA Synthesis Kit (NEB) following the manufacturer's protocol, and synthesized RNA was isolated using the Monarch RNA Cleanup Kit (NEB). Purified RNA samples were stored at −80 °C prior to use.

#### Synthesis of peptide libraries

Synthesis was conducted using the standard Fmoc peptide synthesis methodology using the SPENSER automated synthesizer (IOCB Prague, Czech Republic). The synthesis of the libraries was conducted on 384-well filter plates (Cytiva 5072-N) with a reaction scale of 1410 nmol per well. Incubations were carried out using an apparatus that alternated between N2 overpressure (0.05 atm) and suction (−0.75 atm) to mix the well contents, with a period of 5 s between 200 ms suction pulses. To start the synthesis, Fmoc-protected amino acids were weighed, combined, and then dissolved to form a 0.3 M solution with 0.375 M OxymaPure in dimethylformamide (DMF) to form an isokinetic mixture.
Deprotection was performed by sequential addition of 6 × 40 µL of 20% piperidine in DMF, with each addition incubated for 30 min. Wells were washed using 35, 80, and 6 × 35 µL of DMF. Supernatants were removed via suction applied for 1 min. Coupling was carried out using 2 × 45 µL of isokinetic mixture, and 375 mM N,N'-diisopropylcarbodiimide in DMF and incubated for 3 h each time, followed by washing, deprotection and washing. After the final deprotection cycle, the resin was thoroughly rinsed with DMF and all liquid contents were removed via suction applied for 1 min. After that, resin was taken out of the 384-well filter plate via centrifugation at 4000 × *g* for 5 minutes and combined. The dried, combined resin underwent treatment either with fluorescent labelling mixture or was cleaved directly.
Fluorescent labelling was accomplished on dried resin, to which 5(6)-carboxytetramethylrhodamine was added as 0.3 M solution in DMF (3 eq.) with 0.375 M OxymaPure as additive and 0.375 M N,N'-diisopropylcarbodiimide as coupling reagent.
Cleavage was accomplished with 10 ml of Mixture K (trifluoroacetic acid–water–triisopropylsilane, 95:2.5:2.5, v/v) per gram of resin for 2 h. The resin was then filtered out, and the resulting peptide solution was precipitated with diethyl ether (20 ml/1 ml Mixture K) and centrifuged at 4000 × *g* for 5 minutes. The resulting pellet was suspended in 1 mM HCl and subjected to lyophilization. Before conducting subsequent experiments, all peptide libraries were further lyophilized thrice from 1 mM HCl overnight, establishing Cl- as the counterion.

To perform Quality Control of peptide libraries, the molecular weight distributions of combinatorial peptide libraries were confirmed by mass spectrometry using UltrafleXtreme MALDI-TOF/TOF mass spectrometer (Bruker Daltonics, Bremen, Germany) according to the standard procedure. Prior to the amino acid analysis, the library samples were hydrolyzed in 6 M hydrochloric acid at 110 °C for 20 h and the hydrolyzate was evaporated and reconstituted with loading buffer (pH 2.20) containing an internal standard. Amino acid analysis was performed on a Biochrom 30+ analyzer (Biochrom, Ltd., UK).
The presence of 2,4-diaminobutyric acid was confirmed by fluorescamine assay. For the assay, peptide library solution (75 µL) in PBS buffer was incubated with 25 µL of fresh stock of 3 mg/mL fluorescamine in DMSO at room temperature for 1h in a 96-well plate (Greiner 650209). Fluorescence intensity (λEx = 365 nm, λEm = 470 nm) was recorded using a Tecan Spark plate reader.
Primary amine concentrations were determined by linear interpolation of measured intensities for peptide H-GTIQPYPFSWGY-NH2 in concentration range 87.5−2.7 µM. Obtained amine concentrations were then divided by the concentration of the analyzed peptide determined by amino acid analysis to calculate the amount of amine equivalents per molecule. Experiments were conducted in duplicate.

**Urea–PAGE Analysis of sPTC Stability**

The chemical stability of 8.4 μM sPTC RNA was evaluated via denaturing urea polyacrylamide gel electrophoresis (urea-PAGE). Samples were formulated in either nuclease-free water or a universal buffer(1): Tris-HCl, 20 mM Bis-Tris, 20 mM sodium acetate) adjusted to pH 4.0, 5.0, 6.0, 7.0, or 8.0. $MgCl_2$ was supplemented to yield $Mg^{2+}$-to-RNA phosphate charge ratios $R_{+/-}$ of 0, 4, 8, 16, and 32. Following incubation at 25 °C for 24 h or 7 d, RNA was resolved on 12% polyacrylamide gels containing 8 M urea. Gels were stained with GelRed (Biotium) and imaged on a G:BOX documentation system (Syngene) utilizing constant exposure parameters to ensure comparability. Densitometric quantification (Fig. S1) was performed in Fiji (ImageJ) by integrating the fluorescence signal within a standardized circular region of interest (ROI) applied uniformly across all bands. Relative RNA stability was determined by normalizing the integrated intensities ($I_i$) to corresponding day 0 controls: $I_i / I_{ref}$ - 1. All experiments were conducted in triplicate.

**sPTC RNA denaturation curves**

Temperature denaturation curves (Fig. S3) of 1 μM sPTC were determined in different conditions: with the addition of either 1 mM $MgCl_2$ and 20 mM Tris - pH 7.6, 1 mM $MgCl_2$ and 4 M urea or 4M urea. The RNA solutions (~600 μL) were prepared in Eppendorf tubes, transferred to standard 1 cm optical path quartz cuvettes and successively covered with mineral oil. Samples were slowly annealed in a standard UV–Vis spectrophotometer at 1°C/min from 20 °C to 95 °C, then left for 2 minutes at 95°C and then cooled down at the same rate. Absorbance spectra in the 210-320 nm range were collected and the raw signals around the 260 nm peak were then averaged. Baselines were corrected by fitting with an exponential drift the average signal from the spectral region in the [210,220] nm range. Data refer to cooling temperature ramps.

**Temperature-dependent UV absorbance of sPTC in the presence of $Mg^{2+}$**

Thermal melting profiles of sPTC (Fig. S5) were recorded by monitoring absorbance at 260 nm using a Specord 50 Plus UV–Vis spectrophotometer. Measurements were performed in quartz cuvettes using 230 μL samples, containing 2 μM sPTC and $MgCl_2$ at $R_{+/-}$ = 16, in either $H_2O$ or presence of universal buffer at pH 4. Temperature was increased at a constant rate of 0.5 °C/min and the absorbance at 260 nm was simultaneously monitored throughout the temperature ramp.

**Microscopy Analysis of $Mg^{2+}$-Induced Phase Separation**

Phase separation of 8.4 μM sPTC RNA was evaluated in the presence of $Mg^{2+}$ at charge ratios $R_{+/-}$ of 0, 4, 8, 16, and 32. Samples were prepared in either nuclease-free water or universal buffers (pH 4.0, 5.0, 6.0, 7.0, and 8.0) to assess pH dependencies. For thermal screening, aqueous samples were heated from room temperature (RT) to target temperatures (25, 40, 55, 70, 85, or 95 °C) at a rate of 0.1 °C $s^{-1}$, held for 10 s, and cooled to RT at 0.1°C $s^{-1}$. For pH screening, buffered samples were subjected to an identical thermal cycle to a peak temperature of 95 °C. Following thermal cycling, all samples were incubated at RT and analyzed at 24 h and 7 d intervals. All microscopy measurements were performed in 96-well flat-bottom Costar plates (10 μL sample volume) using a Leica DMi8 inverted microscope equipped with a 40× objective. Bright-field and Differential Interference Contrast (DIC) optics were employed to characterize condensates formation, morphology and temporal stability (Fig. S6).

**Coacervation**

RNA solutions were prepared in RNase free water in small aliquots (2-4 μl) and at various concentrations, and they were stored at −80 °C. Lyophilized peptides and peptide libraries were suspended in RNase free water to typical concentrations of 1-5 mM and stored at −20°C. Phase separation diagrams were estimated as a function of the $R_{+/-}$ ratio and of the molar charge concentration (in units of the elementary charge) Q of the mixtures defined as:

$$R_{+/-} = \frac{C_{pep}\, z_{pep}}{C_{RNA}\, z_{RNA}}$$

$$Q = C_{pep}\, z_{pep} + C_{RNA}\, z_{RNA}$$

where $C_{pep}$ and $C_{RNA}$ are the molar concentrations of peptides/peptide libraries and rRNA, respectively. The $R_{+/-}$ ratio represents the stoichiometric charge ratio of the molecules in solution (positive over negative charges), while Q denotes the molar concentration of charged units. Regarding sPTC, $z_{RNA} = 136$, the length of the sequence, while individual peptides net charges $z_{pep}$ are taken from the pH 7.5 estimates given by the Peptide Property Calculator by NovoPro (https://www.novoprolabs.com/tools/calc_peptide_property), and converted in integers: pL2 WT = +4, pL22 WT = +7. For early variants, fully protonated Dab and Dpr are assumed: pL2 UO = +6, pL22 UO = +7. For peptide libraries, $z_{pep}$ is the charge of a representative sequence given the nominal U(R), O(K), E, D percentages (Fig. S8), namely 8E+UO(KR)ED = 0, 8E+UO(KR) = +5, ribUO(KR) = +7, while $C_{pep}$ is the average molarity of the sequences pool based on the nominal amino acids distribution (Fig. S8).

Mixtures of sPTC and peptides/peptide libraries were prepared at room temperature at various working concentrations. Samples were prepared in a total volume of 10 µL in 200 µL Eppendorf tubes, mixing the components in this given order: 1 µl of 200 mM Tris, 1 µl of 10 mM $MgCl_2$, 4 µl of RNA and 4 µl of peptide/peptide library (all samples contained therefore 20 mM Tris pH 7.5 and 1 mM $MgCl_2$). Before adding peptides, the RNA was let equilibrate for 10-30 minutes. After the final addition of peptides the solution was mixed by gentle pipetting.

Phase diagrams were characterized through bright field microscopy. Before imaging, samples were transferred in plastic multiwells (Amine Treated 96 well plates) treated with a Bovine Serum Albumin (BSA, Sigma-Aldrich) 3% in mass solution (~80µL of solution was added for about 20 minutes, then removed and washed twice with ~100µL RNase free water). To avoid contamination and evaporation of the samples, each well was covered with ~100µL mineral oil (M5904, Sigma-Aldrich) and a plastic PCR film was applied on the plate. Samples were imaged using a Nikon TE200 microscope, equipped with a Nikon DS-5M CCDcamera and temperature-controlled chamber, which was set at 25 °C. Before imaging, coacervate droplets were left sedimenting for about 2 hours. Images were taken at different magnifications (10×, 20×, 50×) to characterize LLPS onset and condensates size, morphology and stability.

**Confocal Imaging and FRAP measurements**

Fluorescently labelled condensates were prepared for confocal microscopy as for phase diagram characterization, with the addition of: (i) 500 nM TAMRA-labelled peptides/peptide library for sPTC-peptides coacervates; or (ii) 1x SYBR Green fluorescent dye for sPTC-$Mg^{2+}$ condensates. Fluorescently labelled components were added to the mixtures and let equilibrate with sPTC RNA before inducing LLPS by final addition of unlabelled peptides/peptide library (or by performing temperature ramping). Immediately after preparation, samples were sandwiched between two glass BSA-coated coverslips, separated by a silicone gasket of thickness 1.0 mm with a circular aperture of diameter 9 mm (GraceBio-Labs FastWells TM reagent barriers). After the cell was sealed, the samples were allowed to equilibrate for about 2 h at room temperature. The liquidity of the coacervate samples was first assessed by time-lapse imaging and coalescence behaviour inspection. In addition, the molecular mobility within the condensates was estimated using Fluorescence Recovery After Photobleaching (FRAP).

FRAP measurements were performed using a WLL scanning Confocal Microscope (Leica Stellaris 8) equipped with a 63× (NA = 1.40) oil immersion objective (HC PL APO CS2) and a sCMOS camera Leica DFC9000 GTC. The imaging system was operated using the LASX (LEICA) Software. Photobleaching was achieved by 10-20 cycles of a series of wavelengths around the absorbance peak (530 nm for TAMRA and 495 nm for SYBR Green) set at the maximum output. For a given droplet, bleaching is performed on a circular region of interest (ROI) of diameter of the order of 1µm. For each bleached ROI, a sequence of 10 frames is recorded prior to bleaching (typical duration of 2 s) and one of 400 frames immediately after bleaching (typical duration of 1 minute). To monitor photobleaching during the acquisition, only droplets close to at least one other droplet, chosen as a reference, are considered.

For each FRAP measurement, the pre- and post-bleaching raw data series were analyzed without further image pre-processing. A custom *Python* script (relying on the *readlif*, *skimage* and *cv2* packages) was used to perform automated analysis of the .lif data. Briefly, the ROI corresponding to the bleach spot is retrieved from the metadata and used to measure the average pre-bleaching $I_{pre}(t)$ and post-bleaching $I_{post}(t)$ intensities. Using OTSU thresholding (cv2.THRESH_OTSU) and morphological filling (cv2.MORPH_CLOSE) with elliptical kernel (cv2.MORPH_ELLIPSE), droplets are segmented, labeled and the optimal one is chosen as a reference, based on its size and proximity to the bleached one. The average intensity of the reference droplet $I_{ref}(t)$ and the background intensity $I_{bkg}(t)$ are then measured. $I_{ref}(t)$ is used to quantify the unwanted photobleaching due to laser exposure during imaging, while $I_{bkg}(t)$ is treated as noise and its average value is subtracted to $I_{pre}(t)$, $I_{post}(t)$ and $I_{ref}(t)$. The normalized FRAP signal $I_{norm}(t) \in [0,1]$ is then computed as:

$$I^*_{post}(t) = I_{post}(t)\,\frac{\langle I_{ref}(t)\rangle}{I_{ref}(t)} \;\longrightarrow\; I_{norm}(t) = \frac{I^*_{post}(t) - I^*_{post}(0)}{\langle I_{pre}(t)\rangle - I^*_{post}(0)}$$

where $\langle\cdot\rangle$ indicates the time average. In the case of pL22 WT/UO samples (Fig. 3F), the time evolution of $I_{norm}(t)$ is modeled with a simple exponential recovery:

$$I_{norm}(t) = M\left(1 - e^{-t/\tau}\right)$$

where $M$ is the fractional recovery and $\tau = R^2{}_{bleach} \,/\, D_{app}$, with $D_{app}$ being the apparent diffusion coefficient. Fitting the above equation to the experimental normalized FRAP signals allows for the estimation of $D_{app}$ and $M$ as the best-fitting parameters.

**Simulations of bare RNA**

Systems were simulated at four temperatures: 300, 350, 400, and 450 K. Each system was equilibrated gradually through several steps. First, the potential energy was minimized using the steepest descent algorithm over approximately 13,000 steps. The system was then heated from 10 K to the target temperature over a 100-ps constant-volume simulation using the v-rescale thermostat, followed by a 1-ns constant-pressure simulation with a reference pressure of 1 bar using the Berendsen barostat. For each temperature, four independent production trajectories were generated, differing in the initial velocity sets assigned during equilibration. Each production simulation spanned 500 ns and employed the v-rescale thermostat (300 K) together with the Parrinello-Rahman barostat (1 bar).

Equations of motion were integrated using the leap-frog algorithm with time steps of 1 fs and 2 fs for the equilibration and production stages, respectively. Covalent bonds were constrained to their equilibrium lengths using the P-LINCS algorithm. Electrostatic interactions were treated with the particle mesh Ewald method using a direct-space cutoff of 1.0 nm, while van der Waals interactions were described by the Lennard-Jones potential with a cutoff of 1.0 nm. All simulations were performed using GROMACS 2025.

The radius of gyration was calculated using the *gmx gyrate* tool from the GROMACS package. Intramolecular hydrogen bonds within the RNA were identified using *gmx hbond*, with the default interatomic distance threshold of 0.35 nm and angular threshold of 30°. The number of $Mg^{2+}$ ions coordinated to the RNA was determined using a custom Python script employing the MDAnalysis library, with a $Mg^{2+}$ ion considered coordinated if its distance to any RNA atom was below 0.5 nm.

**Simulations of RNA-peptide mixtures**

The RNA-peptide mixture was modelled using a periodic rhombic dodecahedron box with a volume of 5000 $nm^3$ containing one sPTC and 20 peptide molecules, roughly corresponding to a charge ratio $R_{+/-}$ of 1. The pL22 WT and pL22 UO variants were capped with N-methylamide and acetyl groups to eliminate terminal charges. Peptide conformations were sampled from 120-ns simulations of a single isolated peptide in water at 298 K and 1 bar. Twenty copies of the peptide (obtained clustering of the single peptide simulations) were then inserted into the box at random positions and orientations, yielding

10 independent initial configurations. Each box was subsequently solvated with SPC/E water, neutralized with chloride ions, and supplemented with 10 mM excess $MgCl_2$. The system is described by Amber's ff12sb. Non-canonical residues Dab and Dpr were parameterized based on GAFF; for comparison, we also re-parameterized Lys, here denoted as Lym.

Each system was equilibrated using the following protocol. Potential energy was first minimized using the steepest descent algorithm over approximately 10 thousand steps. Temperature was then equilibrated to 298 K using the v-rescale thermostat over a 100-ps simulation, followed by box-size equilibration over a 500-ps simulation using the Berendsen barostat with a reference pressure of 1 bar. Production simulations were then performed at a constant temperature of 298 K and pressure of 1 bar, yielding 300-ns trajectories. For production runs, hydrogen mass repartitioning (factor of 3) was applied to allow a 4-fs integration time step. Equations of motion were integrated using the leap-frog algorithm, and covalent bonds involving hydrogen atoms were constrained using the P-LINCS algorithm. Electrostatic interactions were treated with the particle mesh Ewald method using a direct-space cutoff of 1.0 nm, and the Lennard-Jones potential was truncated at 1.0 nm. All RNA-peptide mixture simulations were performed using GROMACS 2024.

Trajectories were analyzed using a combination of GROMACS analysis tools and custom *Python* scripts. Measured quantities were either averaged between the 10 different replicas and errors estimated as standard error of the mean or cumulated together.

RNA-peptides contacts: Interactions between RNA and peptides are described by counting the amino acids within 2 Å from RNA (Fig. 4B, S15C) using *gmx select* (*-selrpos atom -seltype res*). Per-residue counts are then normalised by the total.

Hydrogen bonds (H-bonds): Hydrogen bonds between peptides and RNA are identified using *gmx hbond* with the default interatomic distance threshold of 0.35 nm and angular threshold of 30° (Fig. 4C, S15A). The number of H-bonds is then normalized by the number of bound peptides in each replica and by the occurrence of each residue in the peptide sequence.

H-bonds lifetime: RNA-Lys/Arg/Dab/Dpr H-bonds are printed frame-wise using the *gmx hbond* with *-pf* option. Using a custom *Python* script, the list of triplets (donor, hydrogen, acceptor) is parsed, and the persistence of every occurring triplet is checked against consecutive frames, producing an histogram of H-bonds lasting at least $t$. The histogram is normalized to produce a pdf $f(t)$, related to the survival function:

$$S(t) = \int_t^{\infty} f(t')\,dt'$$

which is estimated by taking the reverse cumulative sum of the pdf. The $S(t)$ estimates are then interpolated onto a common time axis and averaged between replicas (Fig. 4E). The area under the curve of $S(t)$ gives a robust non-parametric estimate of the survival time (Fig. 4D, S15B).

Buried Surface Area (BSA): The binding interface (equivalently, the BSA) between RNA and peptides (Fig. S12) is evaluated is evaluated using the Solvent Accessible Surface Area (SASA) of sPTC and the peptides:

$$BSA = SASA_{RNA} + SASA_{PROT} - SASA_{RNA+PROT}$$

with RNA referring to sPTC, PROT to the ensemble of 20 peptides and (RNA+PROT) to the union of these two sets. SASAs are computed using *gmx sasa* with the default settings of 0.14 nm solvent probe radius and 24 number of dots per sphere.

Distribution of cations: The occupancy of Arg, Lys and $Mg^{2+}$ ions (Fig. S13) is computed using *gmx select* (options *-of* and *-ofpdb*, *-selrpos atom -seltype res*) with a 0.5 nm cutoff between RNA atoms and cations.

<u>RNA structural analysis:</u> The persistence of the secondary and tertiary structure of the sPTC in mixture with pL22 WT/UO during 200 ns long simulations was assessed by computing the the per-residue Root Mean Square Fluctuation (RMSF) of the RNA (Fig. S14A, C) using *gmx rmsf* (option *-res* for numerical values, options *-q* and *-oq* for PDB structures). The Root Mean Square Deviation (RMSD) of the whole RNA molecule is calculated averaging the per-residue RMSF values. Additionally, the secondary structure matrix of the RNA (Fig. S14B) is estimated by computing the average minimum distance of every base pair by means of *gmx mdmat* (cutoff of 1 nm, *-nlevels 50*) on the last 50 ns of the trajectories.

**Amino acid Analysis**

Samples for amino acid analysis were placed in glass test tubes, evaporated, mixed with 0.1 mL constant-boiling hydrochloric acid (analytical grade, 6 M), vacuum-sealed, and hydrolyzed at 110°C for 20 hours. After hydrolysis, we evaporated the samples, mixed them with 100 uL loading buffer (pH 2.2, 0,2 M sodium citrate), and injected. Analyses were performed using a Biochrom 30+ amino acid analyzer (Biochrom, UK), where amino acids separate on a column of cation-exchange resin, using three sodium citrate buffers with different pH values and molarities (pH 3.20, 0.2 M; pH 4.25, 0.2 M; and pH 8.60, 0.5 M sodium citrate-based) and one regeneration solution (0.4 M sodium hydroxide) for stepwise elution, also involving a temperature gradient. The eluent combines with a proprietary ninhydrin reagent in a thermostated capillary reactor. Colored products register on a two-channel photometer (550 nm for primary and 440 nm for secondary amino acids). Quantification entails a three-point calibration with a mixed standard (AAS18, Sigma-Aldrich, supplemented with the two dibasic amino acids) (DataSet1).

### Supplementary Figures

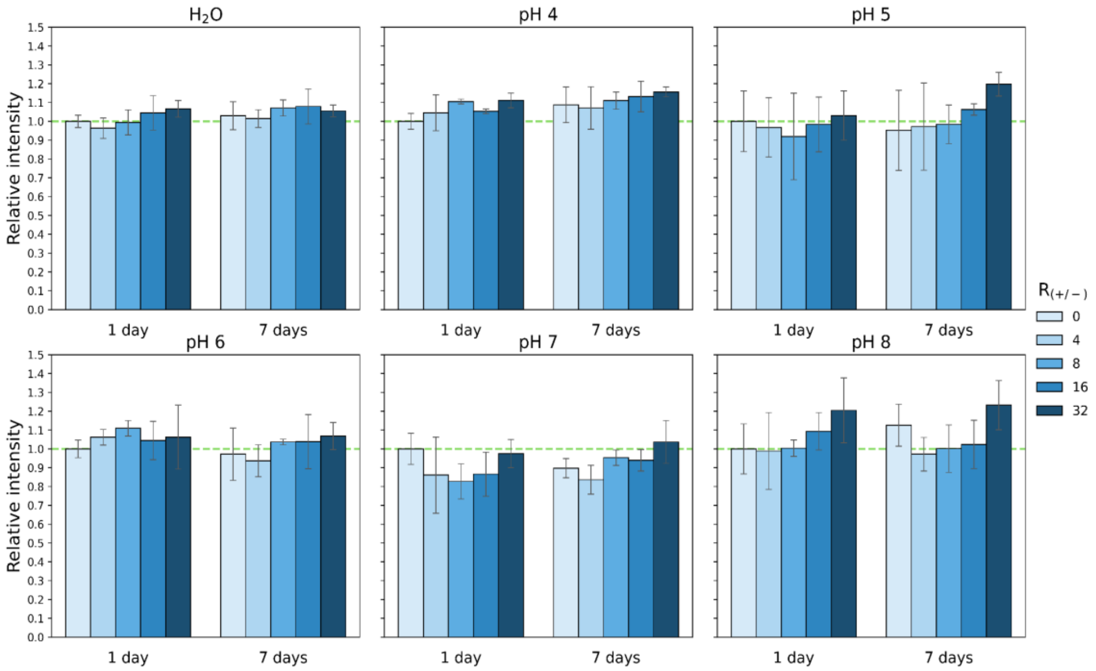


**Figure S1.** Stability characterization of sPTC**.** Triplicate urea-PAGE gels were prepared, visualized, and analyzed for sPTC at pH 4–8 and in pure water. The first five lanes in each gel correspond to samples incubated for 24 h at charge ratios $R_{+/-}$ = 0, 4, 8, 16, and 32, from left to right, with different shades of blue indicating the respective charge ratios. The remaining five lanes correspond to the same charge ratios after 7 days of incubation. Barplots show the mean intensity ± SD across three independent gels.

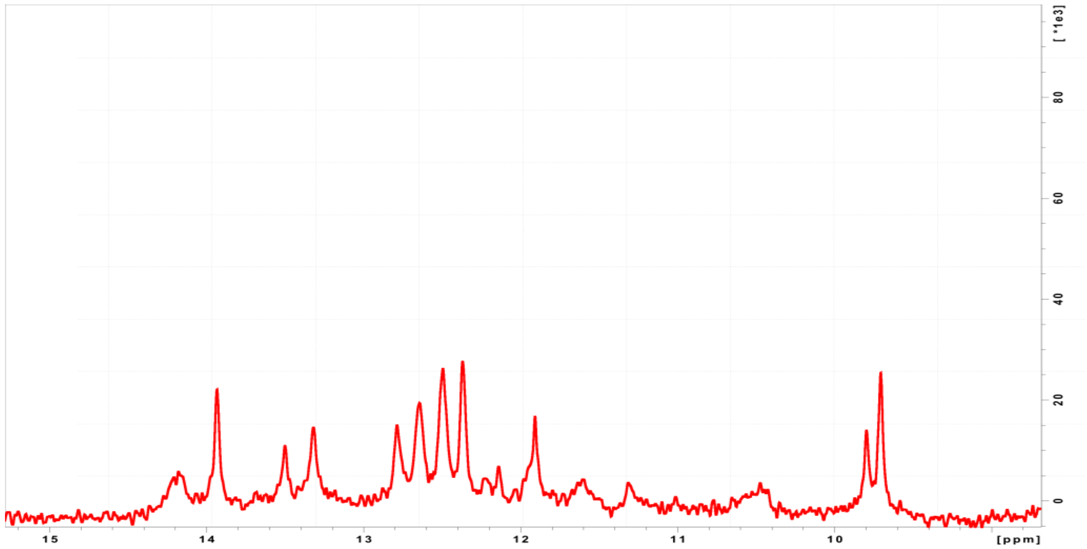


**Figure S2.** 1D NMR spectra of sPTC. One-dimensional ^1H NMR spectrum of sPTC recorded in $H_2O$ at 25 °C. The spectrum shows the 9–15 ppm region for a 60 µM sPTC sample.

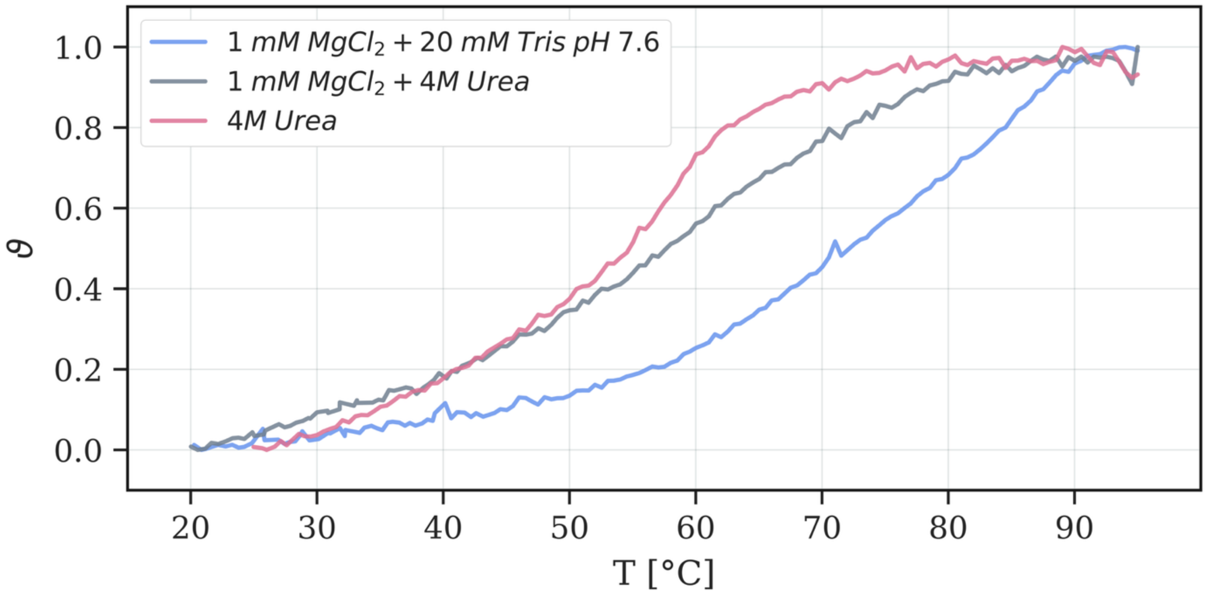

**Figure S3.** Temperature denaturation curves of sPTC. Normalized absorbance melting curves of sPTC at 1 µM in RNase free water with the addition of either 1 mM $MgCl_2$ and 20 mM Tris - pH 7.6 (blue curve), 1 mM $MgCl_2$ and 4 M urea (gray curve) or 4M urea (red curve).

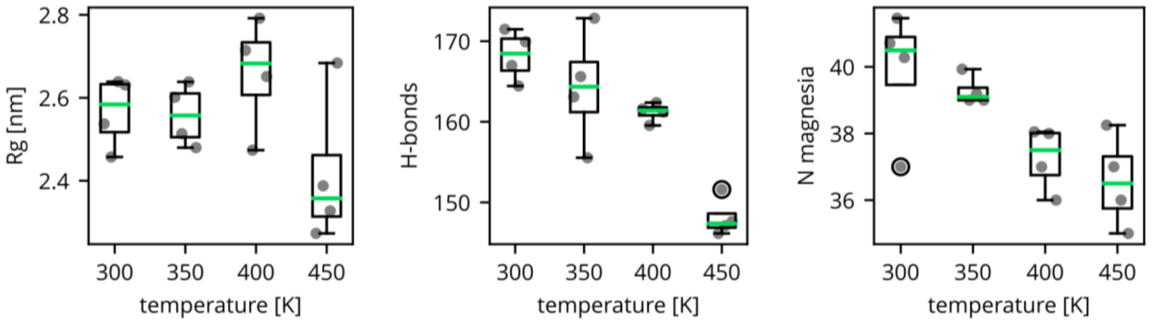


**Figure S4.** Molecular dynamics simulations of sPTC stability. The radius of gyration (Rg), the number of hydrogen bonds (H-bonds) within the sPTC RNA, and the number of $Mg^{2+}$ ions bound to the sPTC ($Mg^{2+}$ distance from sPTC < 0.55 nm) are shown. The dots represent averages over the final portions (400-500 ns) of four independent trajectories. The green lines are medians of these averages, the boxes represent the interquartile ranges, and the whiskers extend from the box to the farthest data point lying within 1.5x the IQR from the box. The fliers beyond this distance are represented by black circles.

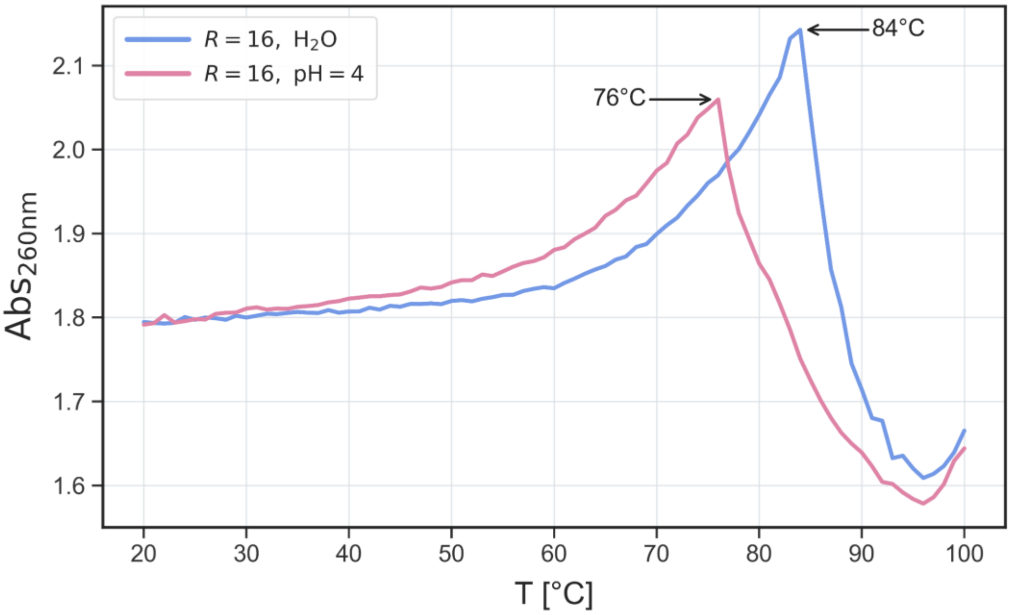


**Figure S5.** Temperature-dependent UV absorbance of sPTC in the presence of $Mg^{2+}$. UV absorbance of sPTC (2 µM) monitored at 260 nm in the presence of $Mg^{2+}$ at $R_{+/-}$ = 16. Signals were recorded from 20 to 100 °C either in $H_2O$ (*blue*) or in universal buffer at pH 4 (*red*). The temperatures marked by arrows correspond to the onset of turbidity, signalling phase separation.

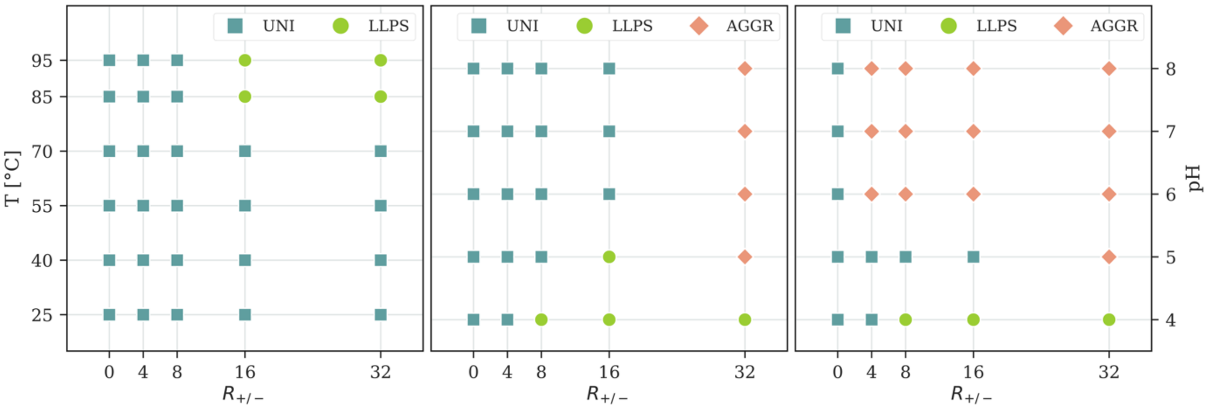


**Figure S6.** $Mg^{2+}$-sPTC phase behaviour as a function of temperature, pH, and $R_{+/-}$. (*left*) Experimental phase diagram of sPTC as a function of temperature and $R_{+/-}$ after 24 h. The corresponding phase diagram after 7 days is not shown because it displayed no detectable difference. (*middle*) Experimental phase diagram as a function of pH and $R_{+/-}$ after 24 h. (*right*) Experimental phase diagram as a function of pH and $R_{+/-}$ after 7 days. Blue squares indicate mixed states (UNI), red diamonds indicate the presence of aggregates (AGGR) and green circles indicate the formation of liquid-like droplets (LLPS).

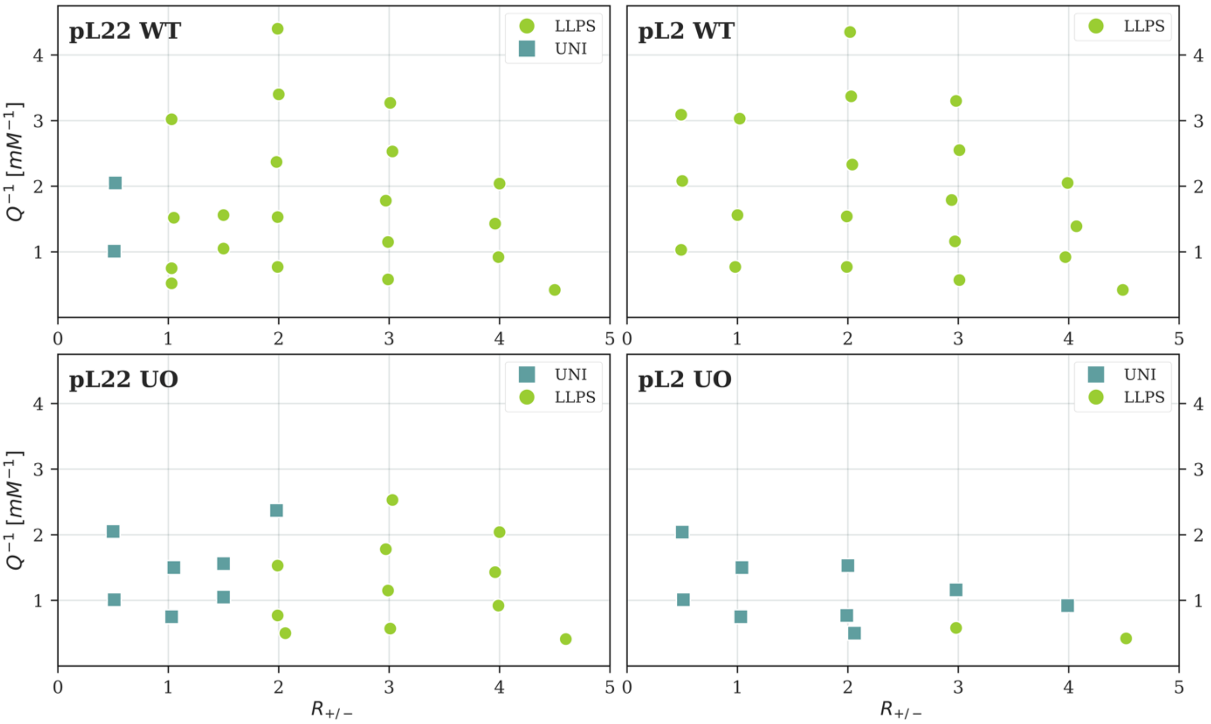


**Figure S7.** Experimental phase diagrams of sPTC with ribosomal peptides and their prebiotic variants. Phase diagrams of sPTC with pL22 WT *(top left*), pL2 WT (*top right*), pL22 UO (*bottom left*) and pL2 UO (*bottom right*) as a function of the stoichiometric charge ratio $R_{+/-}$ and of the inverse of the molar concentration of charged units Q. Blue squares indicate mixed states (UNI) and green circles indicate the formation of liquid-like droplets (LLPS).

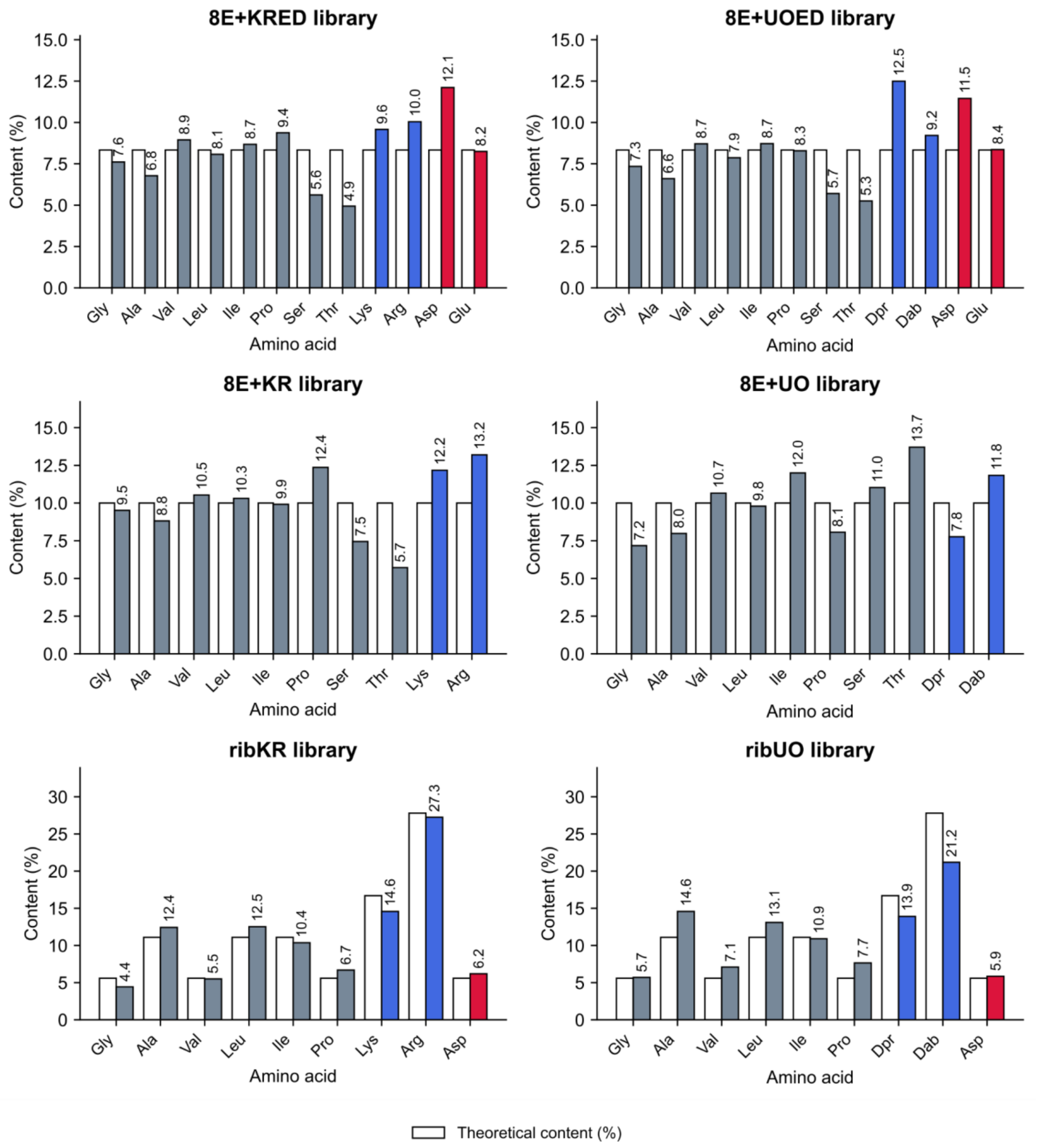


**Figure S8.** Amino acid composition of peptide libraries assessed by HPLC amino acid analysis. Nominal content (white bars) represents equimolar (8E+KR/KRED; 8E+UO/UOED) or rpL22 (ribKR/UO) distribution of amino acids and experimental distributions (coloured bars) show integrated ion intensities from HPLC chromatograms (Dataset1). The 8E+KR/KRED and 8E+UO/UOED libraries are 25 amino acids long, whereas the ribKR/UO libraries are 18 amino acids long.

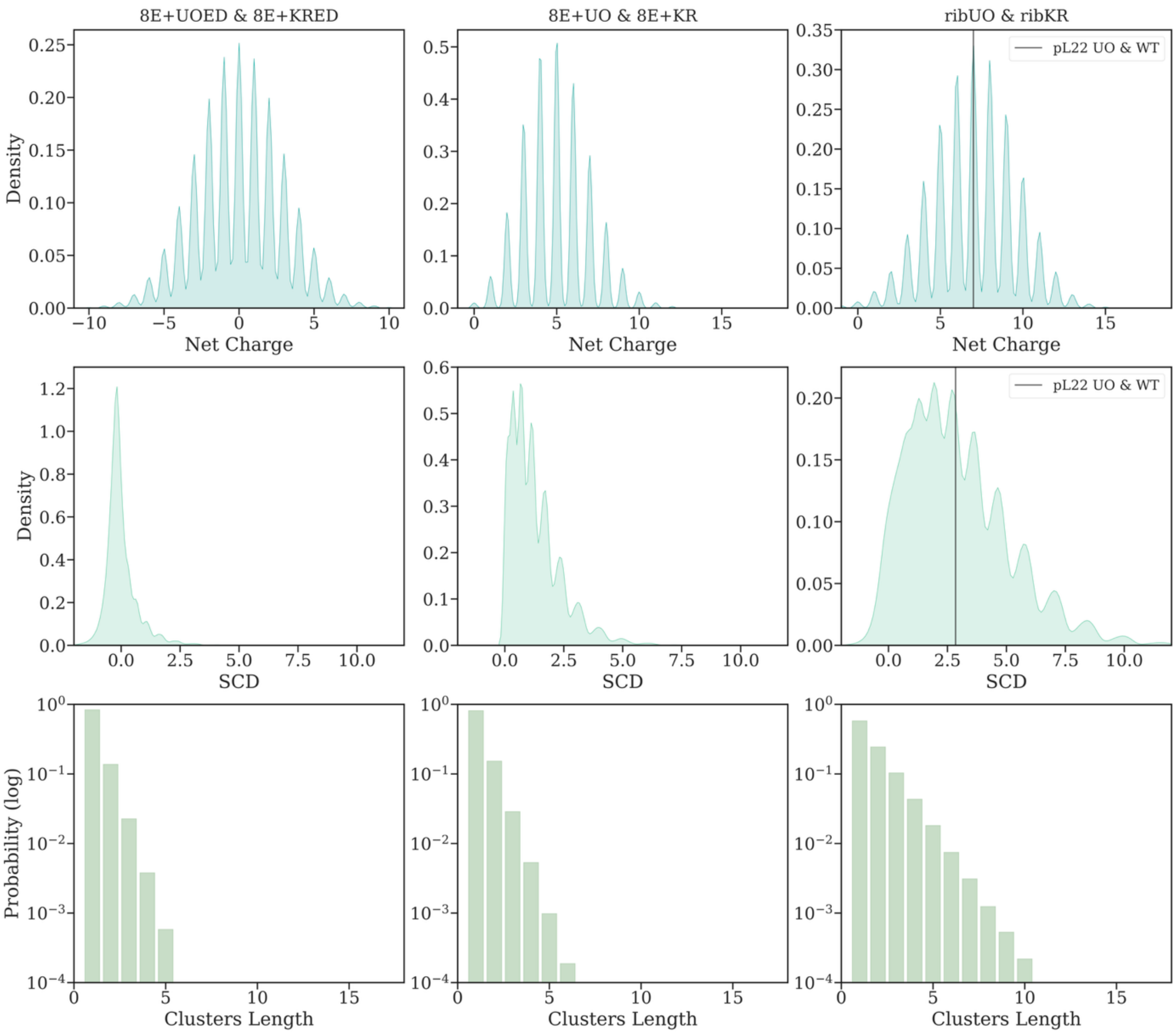


**Figure S9.** Estimated distributions of net charge, SCD and positive charge clusters in peptide libraries. (*top row*) Net charge probability density functions of 4x10^5 peptide sequences randomly generated according to the theoretical amino acid composition of the 8E+UOED and 8E+KRED libraries (*left*), the 8E+UO and 8E+KR libraries (*middle*) or the ribUO or ribKR libraries (*right*). (*middle row*) Same as the top row, but for the Sequence Charge Decoration (SCD) metric defined as:

$$SCD(s) = \frac{1}{N} \sum_{m=1}^{N} \sum_{n=0}^{m-1} q_m q_n (m-n)^{1/2}$$

(*bottom row*) Same as the top row, but for the cluster size of positive charges defined as the length of consecutive positive charge stretches.

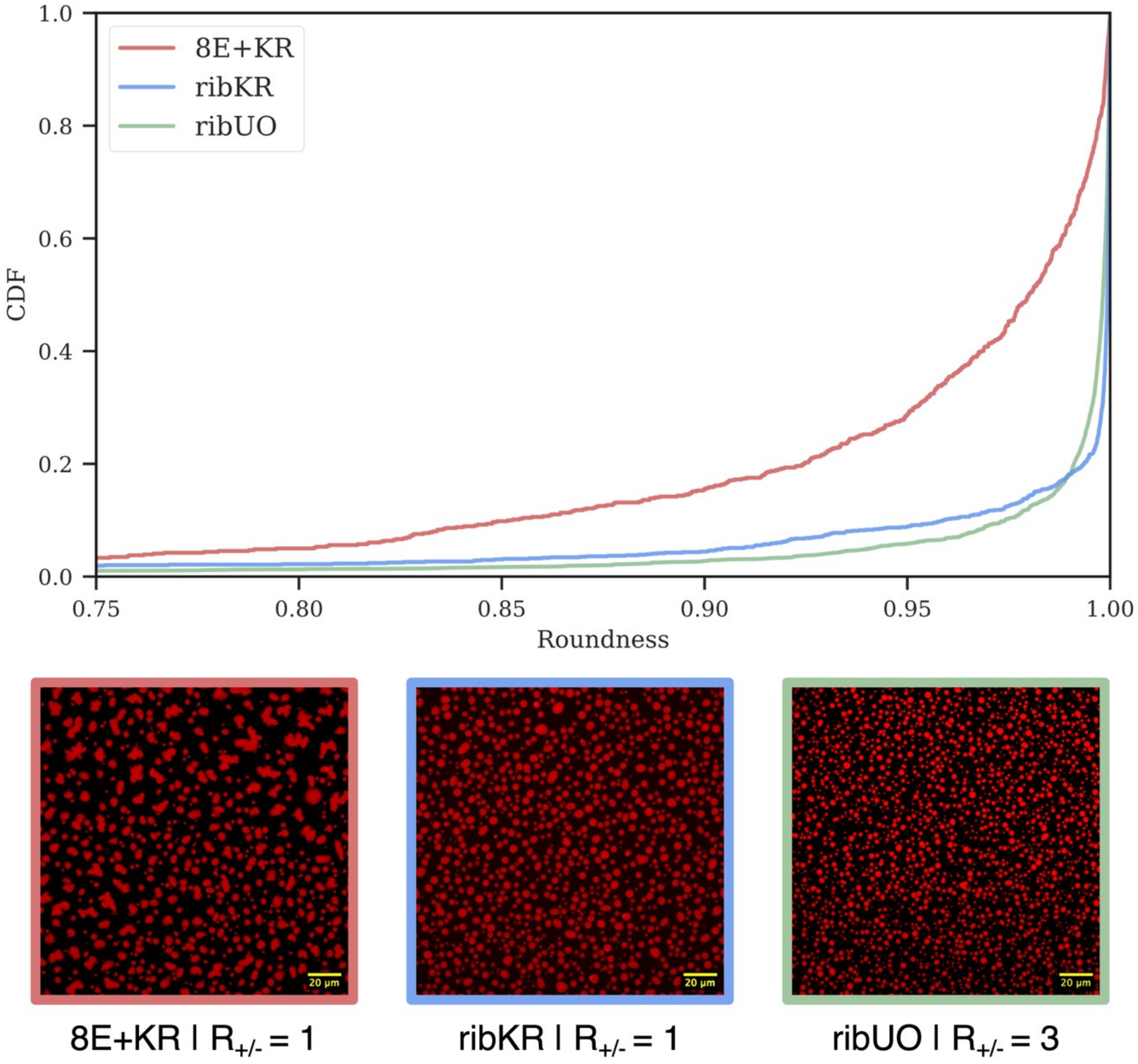


**Figure S10.** Distribution of droplet roundness in sPTC condensates. Cumulative distribution functions (CDFs) of the roundness of sPTC condensates with 8E+KR (*red*), ribKR (*blue*) or ribUO libraries (*green*), with Q ~ 1.3 mM. CDFs are estimated from image segmentation of z maximum projected confocal images (shown below, field of view of 185 um X 185 um). Segmentation was achieved using *Cellpose* v2.2.2 (CP model, flow_threshold 0.4, cellprob_threshold 0.0). Roundness is defined as the ratio of perimeter to area of an equivalent ellipse.

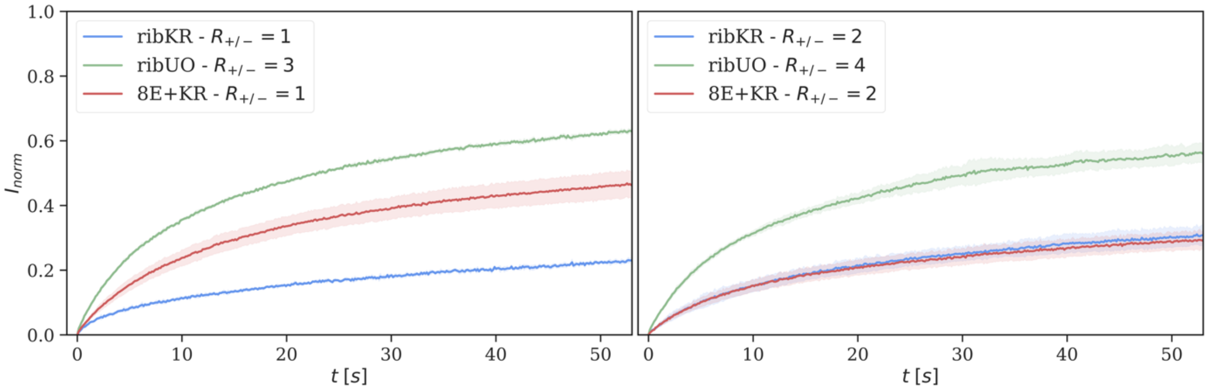


**Figure S11**. FRAP of sPTC condensates with peptide libraries. Averages and standard deviations of normalized FRAP signals from 8E+KR (*red*), ribKR (*blue*) or ribUO (*green*) condensates (n=6). Condensates on the left panel are prepared at Q ~ 1.3 mM, while on the right Q ~ 0.86 mM.

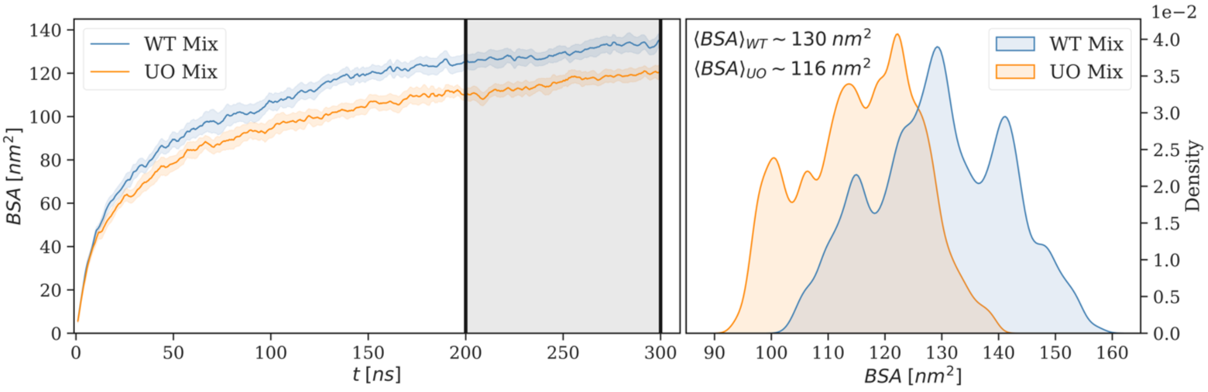


**Figure S12.** Binding interface of pL22 WT *vs* pL22 UO with sPTC from MD simulations. Buried Surface Area (BSA) between the sPTC and pL22 WT (*blue*) / pL22 UO (*orange*) ensembles. On the left, the average and standard error of 10 independent replicas of the BSA time trace shows peptides binding during the first 50 ns. On the right, the cumulative distribution of the BSA during the last 100 ns shows significantly higher values for WT peptides with respect to UO variants.

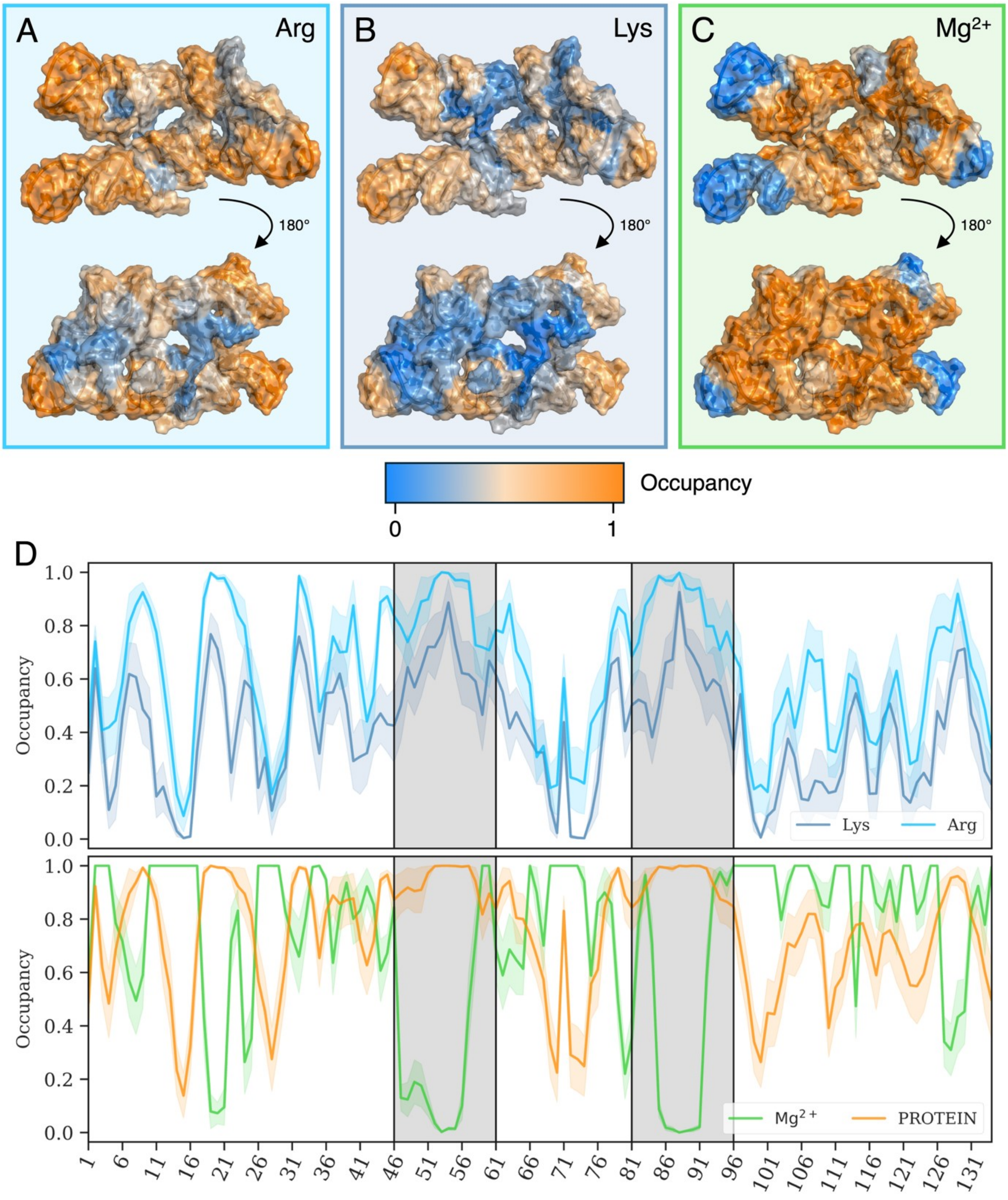


**Figure S13.** Spatial distribution of positive moieties binding to sPTC from MD simulations**.** Occupancy values of Arg, Lys and $Mg^{2+}$ around 0.5 nm of the sPTC during a 100 ns long simulation with pL22 WT peptides, starting from an equilibrated bound state. (A) Occupancy of Arg residues mapped on the sPTC structure. The view on the bottom shows the "back" of the RNA structure shown on top. (B) Same as panel A but for Lys residues. (C) Same as panel A but for $Mg^{2+}$ ions. (D) Occupancy values along the sPTC RNA sequence for Lys (*dark blue*), Arg (*light blue*), $Mg^{2+}$ ions (*green*) and all the pL22 residues (*PROTEIN, orange*). Lines show average values over 10 independent replicas, while shaded areas the respective standard errors. The shaded black boxes mark regions devoid of $Mg^{2+}$ bound ions but enriched in Lys and Arg contacts.

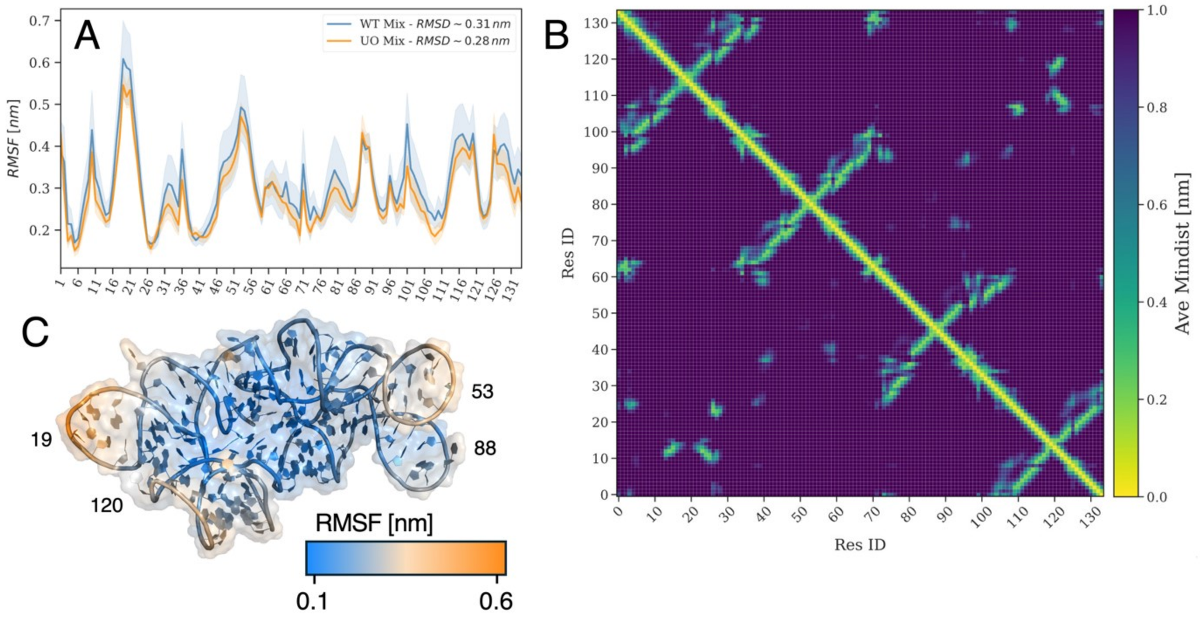


**Figure S14.** Stability of sPTC secondary and tertiary structures from MD simulations. Structural preservation of the sPTC in mixture with pL22 WT or pL22 UO during 200 ns long simulations. (A) per-residue averaged RMSF along the sPTC RNA sequence for mixtures with pL22 WT (*blue*) and pL22 UO (*orange*). Lines represent averages over 10 independent replicas, while shaded areas the respective standard errors. The RMSD of the sPTC is shown in the legend. (B) Secondary structure matrix of the sPTC RNA in mixture with pL22 WT. The average of 10 independent replicas during the last 50 ns is shown. (C) Representative sPTC structure overlaid with the RMSF values shown in panel A. The numbers correspond to the same sequence IDs as in panels A and B.

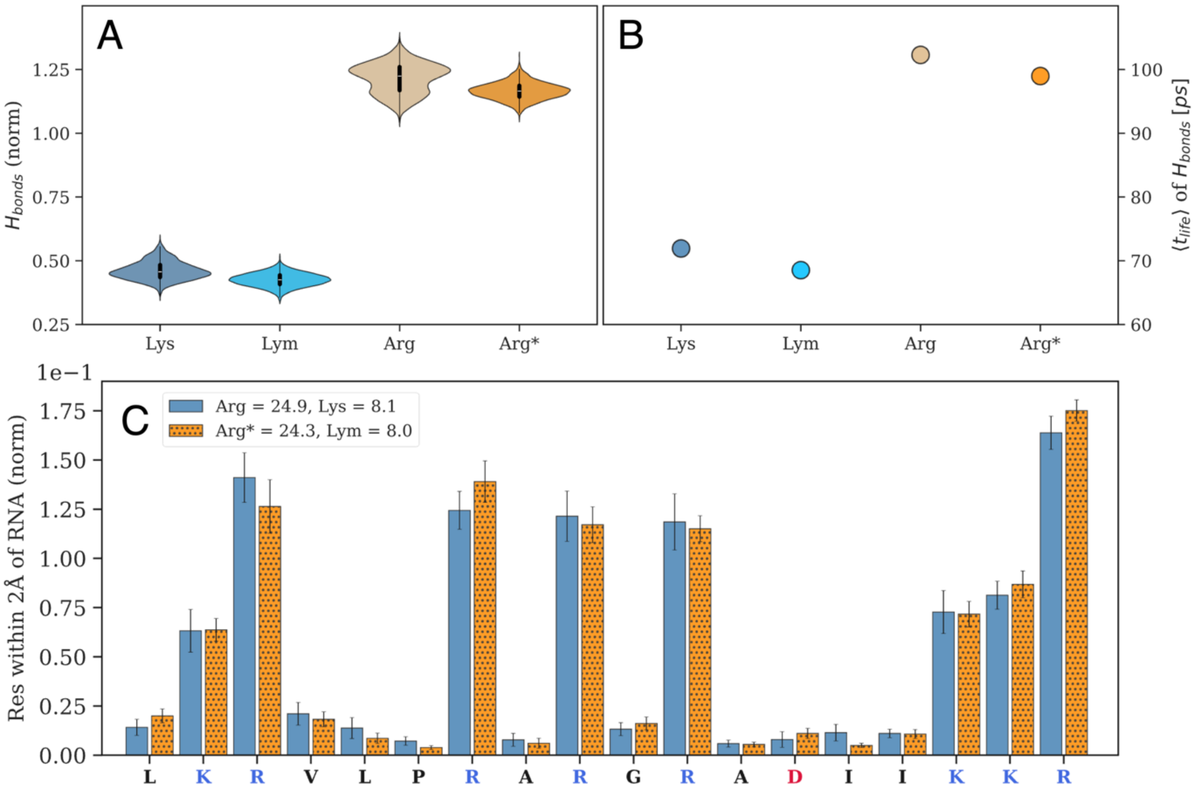


**Figure S15**. Dependence of the results on side-chain parameterization. Lys is compared with its re-parameterized variant Lym, whose parameters were assigned consistently with those of Dab and Dpr. Arg and Arg* have identical parameters, the two labels indicating simulations performed alongside Lys and Lym, respectively. For the sPTC–pL22 WT mixture, the 200 ns MD simulations show no appreciable differences in the normalized number of hydrogen bonds (A), their mean lifetime (B), or the normalized number of contacts per residue along the peptide sequence (C). These measures correspond to those shown in Figure 3B–D.

**Supplementary Tables**

| Name | Sequence |
|---|---|
| sPTC_F | GCGTAATACGACTCACTATAGGAAGACCCCGTGGAGCTCTTCGGAGTTACCCCG GGGATAACAGGCTGATCTCTTCGGAGGTTTGGCAC |
| sPTC_R | GACCGAACTGTCTCACGACGTTCTGAACCCAGCTCGCGTGCCGCCGAAGCGACG AGCCGACATCGAGGTGCCAAACCTCCGAAGAG |
| sPTC | GGAAGACCCCGTGGAGCTCTTCGGAGTTACCCCGGGGATAACAGGCTGATCTCT TCGGAGGTTTGGCACCTCGATGTCGGCTCGTCGCTTCGGCGGCACGCGAGCTG GGTTCAGAACGTCGTGAGACAGTTCGGTC |

**Table S1.** DNA oligos used for preparation of sPTC RNA constructs and sPTC sequence.

| Peptide | Sequence |
|---|---|
| pL2 | H-GRRPHVRGAAMNPVDHPHGGGEGRAPRGR-$NH_2$ |
| pL2 Fluo | 5(6)-TAMRA-GRRPHVRGAAMNPVDHPHGGGEGRAPRGR-$NH_2$ |
| pL2 UO | H-GUUPOVUGAASEOVDOPOGGGEGUAPUGU-$NH_2$ |
| pL2 UO Fluo | 5(6)-TAMRA-GUUPOVUGAASEOVDOPOGGGEGUAPUGU-$NH_2$ |
| pL22 | H-LKRVLPRARGRADIIKKR-$NH_2$ |
| pL22 Fluo | 5(6)-TAMRA-LKRVLPRARGRADIIKKR-$NH_2$ |
| pL22 UO | H-LOUVLPUAUGUADIIOOU-$NH_2$ |
| pL22 UO Fluo | 5(6)-TAMRA-LOUVLPUAUGUADIIOOU-$NH_2$ |

**Table S2.** Sequences of peptides used in this study.
